# Lineage tracing of *Shh+* floor plate cells and dynamics of dorsal-ventral gene expression in the regenerating axolotl spinal cord

**DOI:** 10.1101/2024.06.14.599012

**Authors:** Laura I. Arbanas, Emanuel Cura Costa, Osvaldo Chara, Leo Otsuki, Elly M. Tanaka

## Abstract

Both development and regeneration depend on signalling centres, which are sources of locally secreted tissue-patterning molecules. As many signalling centres are decommissioned before the end of embryogenesis, a fundamental question is how signalling centres can be re-induced later in life to promote regeneration after injury. Here, we use the axolotl salamander model (*Ambystoma mexicanum*) to address how the floor plate is assembled for spinal cord regeneration. The floor plate is an archetypal vertebrate signalling centre that secretes *Shh* ligand and patterns neural progenitor cells during embryogenesis. Unlike mammals, axolotls continue to express floor plate genes (including *Shh*) and downstream dorsal-ventral patterning genes in their spinal cord throughout life, including at steady state. The parsimonious hypothesis that *Shh*+ cells give rise to functional floor plate cells for regeneration had not been tested. Using HCR *in situ* hybridisation and mathematical modelling, we first quantitated the behaviours of dorsal-ventral spinal cord domains, identifying significant increases in gene expression level and floor plate size during regeneration. Next, we established a transgenic axolotl to specifically label and fate map *Shh*+ cells *in vivo*. We found that labelled *Shh+* cells gave rise to regeneration floor plate, and not to other neural progenitor domains, after tail amputation. Thus, despite changes in domain size and downstream patterning gene expression, *Shh*+ cells retain their floor plate identity during regeneration, acting as a stable cellular source for this regeneration signalling centre in the axolotl spinal cord.

## Introduction

Understanding how to regenerate the spinal cord after injury is a central question in regenerative research. Regenerative species such as axolotls *(Ambystoma mexicanum)* and zebrafish *(Danio rerio)* have revealed that important contributors to spinal cord regeneration are resident neural progenitor cells (also known as neural stem cells, ependymal glial cells or ependymoglial radial cells). These progenitors, which line the central canal of the spinal cord, can replace tissue lost or damaged in several injury paradigms in salamanders and zebrafish, such as crush injury (Hui et al., 2010; Thygesen et al., 2019; Walker et al., 2023), transection injury (Becker et al., 1997; Piatt, 1955) or a full tail amputation (Egar and Singer, 1972). For example, amputation of the axolotl spinal cord recruits neural progenitors residing within a ∼800 μm zone to switch to fast, proliferative cell divisions (Albors et al., 2015; Cura Costa et al., 2021; Mchedlishvili et al., 2007; Rost et al., 2016), generating a neuroepithelial tube that differentiates into a functional spinal cord.

The embryonic origin of spinal cord neural progenitors, and their patterning, is well understood in mouse *(Mus musculus)* and chicken *(Gallus gallus)*. In these species, two major signalling centres in the developing neural plate generate opposing morphogen gradients that provide dorsal-ventral positional information (reviewed in (Sagner and Briscoe, 2019)). The dorsally located roof plate secretes Bone Morphogenetic Protein (BMP) family members (Liem et al., 1997) and *Wnt*-family proteins (Muroyama et al., 2002), while the ventrally located floor plate secretes Sonic hedgehog (*Shh*) (Echelard et al., 1993). Neural progenitors residing between the two signalling centres receive different concentrations and durations of signalling molecules and acquire distinct dorsal-ventral identities. As a result, neural progenitors express different transcription factors depending on their location (e.g. *Pax7* and *Msx1* dorsally, *Pax6* laterally, *Nkx6.1* ventro-laterally) and generate distinct neuron subtypes (reviewed by (Sagner and Briscoe, 2019)). Towards the end of embryogenesis, the mouse spinal cord undergoes molecular changes: *Shh* signalling is extinguished (Cañizares et al., 2020), BMP activity extends ventrally (Cañizares et al., 2020) and the expression of dorsal-ventral patterning genes is altered or diminished (Albors et al., 2023; Ghazale et al., 2019). The adult mouse spinal cord regenerates poorly and resident ependymal cells generate a glial scar after injury (Meletis et al., 2008) instead of restoring neurons and function.

An interesting possibility is that instilling an embryo-like arrangement of roof plate, lateral progenitors and floor plate during adulthood would contribute to reconstitution of developmental mechanisms and replace lost neurons. Regenerative axolotls indeed express roof plate genes (*Msx1, Pax7, BMP2*), lateral patterning genes (*Pax6*) and floor plate genes (*Shh, FoxA2*) in this manner throughout life (Schnapp et al., 2005; Sun et al., 2018). It is tempting to speculate that this arrangement acts as a template to launch appropriate gene cascades and replace missing spinal cord regions after injury. In adult zebrafish, the expression of *shha, nkx6.1, pax6* and *olig2* increases locally following spinal cord transection (Reimer et al., 2009), which could reflect the activation of such gene cascades. In axolotls, *Pax6* and *Pax7* expression decrease 1 day post-tail amputation (Albors et al., 2015), but the later expression dynamics of these, and other, genes have not been quantified. Elucidating the dynamics of dorsal-ventral gene expression after axolotl tail amputation could illuminate mechanisms of spinal cord regeneration conserved across injury paradigms and species.

Here, we quantified the expression of dorsal-ventral patterning genes covering roof plate to floor plate during axolotl spinal cord regeneration. Using mathematical modelling, we extracted gene expression levels and the relative sizes of the dorsal-ventral domains from measurements made with single molecule fluorescent *in situ* hybridisation (smFISH). We found that dorsal-ventral genes increased their expression after amputation, similar to zebrafish transection, but additionally discovered changes in the representation of the dorsal-ventral domains. In particular, we found that the *Shh*+ floor plate almost doubles in size, which is relevant considering that it is an essential signalling centre for regeneration: pharmacological inhibition of *Shh* signalling results in an expanded dorsal domain and blocks axolotl spinal cord outgrowth (Schnapp et al., 2005).

The expansion of the *Shh*+ domain prompted us to address how *Shh+* floor plate cells contribute to the regenerated spinal cord. If continuous *Shh*+ expression reflects a cellular memory and fate restriction, floor plate cells would be expected to produce only floor plate cells during regeneration. However, lineage tracing of single electroporated cells have suggested that axolotl progenitors can change dorsal-ventral identity (Mchedlishvili et al., 2007). Given the expression of ventrally-derived *Shh*, it is plausible that neighbouring progenitors could change between medio-lateral, lateral and dorsal fates but whether *Shh*+ floor plate cells themselves remain lineage-restricted, or can change identities, was not determined (Mchedlishvili et al., 2007). We performed genetic fate mapping of *Shh*+ floor plate cells and found that they exclusively generate more floor plate during axolotl spinal cord regeneration, supporting a fate restriction model.

## Materials and methods

### Axolotl (*Ambystoma mexicanum*) husbandry

All procedures were approved by the Magistrate of Vienna Genetically Modified Organism Office and MA58, City of Vienna, Austria (licences: GZ:51072/2019/16, GZ: MA58-1432587-2022-12, GZ: MA58-1516101-2023-21). Axolotls were raised in Vienna tap water. Axolotl breedings were performed at the IMP by the animal caretaker team. Axolotl sizes are reported in cm, measured from snout to tail tip. Axolotl surgeries, live imaging and tissue harvesting were performed under anaesthesia in 0.015% benzocaine (Sigma-Aldrich E1501, preparation according to (Khattak et al., 2014)). Tail amputations were performed between myotome 8-10 post-cloaca (3-4 cm animals) or halfway between cloaca and tail tip (1.5-2 cm animals (lineage tracings)).

### Axolotl genome and transcriptome reference

Axolotl genome assembly AmexG_v6.0-DD and transcriptome assembly AmexT_v47 (Schloissnig et al., 2021).

### Generation of *Shh* knock-in axolotl

*Shh* knock-in axolotl “*Shh^EGFP-dERCre^*” (tm(*Shh^t/+^:Shh-*P2A*-myr-EGFP-*T2A*-ER^T2^-Cre-ER^T2^*)^Etnka^) was generated by CRISPR/Cas9 and NHEJ-mediated knock-in into the last intron of the *Shh* gene (Fei et al., 2018). De-jellied, 1-cell stage axolotl eggs were injected with injection mix as described in (Khattak et al., 2014), delivered as 2 x 2.5 nl shots. Injection mix recipe: 5 μg Cas9-NLS protein, 4 μg *Shh* gRNA, 0.5 μg *Shh* knock-in cassette, 1 μl Cas9 buffer, diluted to 10 μl in water. Cas9-NLS protein and Cas9 buffer were prepared by the Vienna Biocenter Core Facilities. Axolotls with successful knock-in were recovered by screening for EGFP fluorescence in the posterior limb bud at embryo stage 42-44 using an AXIOzoom V16 microscope (Zeiss). Transgenic individuals were reared to sexual maturity and germline-transmitted offspring were used in all experiments.

*Shh* gRNA was prepared as described in (Fei et al., 2018) by PCR amplification and *in vitro* transcription of the following synthesised oligonucleotides (purchased from Merck):

>*Shh-*gRNA oligo_FWD (target sequence in *Shh* last intron is underlined) GAAATTAATACGACTCACTATAGG<u>CGTACTTCTGGACTTTGG</u>GTTTTAGAGCTAGAAATAGC

>Common-gRNA-REV (Fei et al., 2018)

AAAAGCACCGACTCGGTGCCACTTTTTCAAGTTGATAACGGACTAGCCTTATTTTAACTTGCTATTTCTAGCTCTA AAAC

*Shh* knock-in cassette was assembled in a plasmid by Gibson Assembly, purified using a Plasmid Maxi Kit (Qiagen 12163) and verified by Sanger sequencing prior to egg injection. Knock-in cassette encodes: last *Shh* intron and exon, P2A ‘self-cleaving’ sequence, EGFP fluorescent protein fused with a N-myristoylation sequence, T2A ‘self-cleaving’ sequence, tamoxifen-inducible Cre recombinase, poly-adenylation sequence.

### Other axolotl strains

The following published axolotl strains were used in this study: *d/d* (control strain), tm(*Pax7^t/+^:Pax7-*P2A*-memCherry-*T2A*-ER^T2^-Cre-ER^T2^*)^Etnka^ (Fei et al., 2017), tgSceI(*Caggs:loxP-GFP-dead(Stop)-loxP-mCherry*)^Etnka^ (Kawaguchi et al., 2024), tgSceI(*Caggs:loxP-GFP-loxP-mCherry*)^Etnka^ (Khattak et al., 2013). Nomenclature is according to (Nowoshilow et al., 2021).

### Genetic lineage tracing of *Shh+* cells

*Shh^EGFP-dERCre^* axolotls were mated with memory cassette axolotls of genotype tgSceI(*Caggs:loxP-GFP-dead(Stop)-loxP-mCherry*)^Etnka^ (Kawaguchi et al., 2024). (Kawaguchi et al., 2024)To induce Cre/loxP-mediated recombination, progeny axolotls were treated with 4-hydroxytamoxifen (4-OHT) by bathing, as described in the water-based method of (Khattak et al., 2014). 3 cm axolotls were amputated halfway between cloaca and tail tip and bathed overnight in the dark on days 1, 3 and 5 post-amputation with 2 μM 4-OHT. Successfully recombined individuals were identified by screening for mCherry expression 7 days after the last 4-OHT treatment using an AXIOzoom V16 microscope (Zeiss). For lineage tracing, tails were re-amputated <500 μm posterior to mCherry+ cells. Tail offcuts containing mCherry+ cells were harvested to test the fidelity of labelling. Animals were left to regenerate for 7 days (“short-term tracing”) or 28 days (“long-term tracing”) before harvesting.

### Live imaging

Axolotls were anaesthetised in 0.015% benzocaine (Sigma-Aldrich E1501, preparation according to (Khattak et al., 2014)) and imaged using an AXIOzoom V16 microscope (Zeiss) on the indicated days post-tail amputation. Axolotls were returned to tap water immediately after imaging.

### Tissue harvesting and cryosectioning

Axolotl tails were harvested and fixed overnight at 6°C in 4% paraformaldehyde (PFA), pH 7.4. Fixed samples were washed twice with cold PBS then incubated sequentially with the following solutions overnight at 6°C: (1) 20% sucrose in PBS, (2) 30% sucrose in PBS, then incubated for 3 hours in a 1:1 mix of 30% sucrose/PBS and Tissue-Tek O.C.T. compound (Sakura). Samples were embedded in O.C.T., frozen on dry ice and sectioned immediately (20 mm thickness) or stored at -70 °C. Slides were stored at -20 °C until use.

### Immunofluorescent staining of tissue sections

Slides were brought to room temperature and washed with PBS to remove O.C.T. For DAPI staining only: slides were incubated with 10 μg/ml DAPI solution (Sigma-Aldrich D9542) for 30 mins at room temperature, then washed well with PBS. For immunostaining against PAX6, PAX7 and SOX2: slides were incubated for 1 hour at room temperature in blocking solution (PBS containing 1% BSA (bovine serum albumin) and 0.5% Triton X-100). Slides were incubated overnight at 6°C with primary antibodies diluted in blocking solution. The following day, all slides were washed three times over 3 hours at room temperature with blocking solution. Slides were incubated overnight at 6°C with secondary antibodies diluted in blocking solution. Finally, slides were washed three times with blocking solution and once in PBS before mounting in Abberior MOUNT embedding media for imaging. For immunostaining against SHH: antigen retrieval was necessary. After washing off O.C.T., slides were incubated in undiluted 10X citrate buffer (Dako) for 45 minutes at 65 °C, then washed twice in PBS before blocking and proceeding to antibody staining as for the other antigens. Images were acquired using a spinning disk confocal setup (Olympus IX83 inverted microscope / Yokogawa CSU-W1) and a 40x air objective. Primary antibodies and dilutions used were: anti-PAX6 (rabbit, Biolegend, #901301, 1:200), anti-PAX7 (mouse, DSHB, #Pax7-s, 1:100), anti-SHH (rabbit, Cell Signalling Technologies, #2207S, 1:200), anti-SOX2 (rat, eBioscience, Btjce, 1:200). Primary antibodies were detected using secondary antibodies conjugated to Alexa fluorophores (Thermo Fisher Scientific).

### HCR staining of tissue sections and HCR probe design

Slides were brought to room temperature and washed with PBS to remove O.C.T. HCR *in situ* hybridisation was performed according to the HCR RNA-FISH protocol for fresh/fixed frozen tissue sections (Molecular Instruments, (Choi et al., 2018)), omitting the post-fixation and proteinase K treatment steps. Probe hybridisation buffer, wash buffer, amplification buffer and detection hairpins were purchased from Molecular Instruments. Probe hybridisation was performed at 37 °C for 18 h. Amplification was performed at room temperature for 18-20 h using B1/B2/B5 hairpins conjugated to Alexa-546 or Alexa-647 fluorophores. Following the HCR procedure, slides were incubated with 10 μg/ml DAPI solution (Sigma-Aldrich D9542) for 30 mins at room temperature, then washed well with PBS. Samples were mounted in Abberior MOUNT embedding media for imaging. Images were acquired using a spinning disk confocal setup (Olympus IX83 inverted microscope / Yokogawa CSU-W1) and a 40x air objective.

HCR probes were designed against unique mRNA sequences identified by BLAST alignment against axolotl transcriptome Amex.T_v47 (Schloissnig et al., 2021). Sequences were considered unique if they did not match off-target sequences at more than 36 out of 50 consecutive nucleotides. HCR probes targeting axolotl *Shh* mRNA (Otsuki et al., 2023) were purchased from Molecular Instruments; all other probes (*Nkx6.1, Pax6, Pax7, Msx1, Sox2*) were purchased as oPools at 50 pmol scale from IDT (Integrated DNA Technologies).

### Fluorescence intensity quantifications

Image quantifications were performed using Fiji software (Schindelin et al., 2012). The segmented line tool was used to draw a line trajectory through the region of interest (line thickness: “100” for HCR experiments (measurements were made on maximum intensity projections of 20 μm) and “10” for live dual reporter experiments. The Measure function was used to extract continuous mean gray values for analysis.

### Mathematical modelling of fluorescence data

A detailed description of the mathematical modelling can be found in the Supplementary Information. We used a piecewise constant model in which spatial domains of constant signal are separated by one or more switch points (see Figure S1a). We used a two-step model variant for *Msx1, Pax7, Nkx6.1* and *Shh* (two domains separated by one switch point), and a three-step model variant for *Pax6* (three domains separated by two switch points). We determined domain size by fitting the relevant model to the HCR signal data and inferring the switch point(s). To estimate gene expression, we calculated average HCR signal intensities on either side of the switch point(s) and subtracted background signal from expression signal.

To determine the optimal fits, the mean signal levels for the zones defined by the switch point in the two-step function (or pairs of switch points in the three-step function) were determined. Next, the switch points were systematically varied across the data range. For each potential switch point, the mean signal levels in the resulting zones were calculated. To assess the best-fitting parameter values, the sum of squared errors (SSE) was calculated. See Supplementary Information for details on SSE and all individual fits to the HCR data.

### Statistics and data representation

Statistical analysis and graph plotting were performed using custom Python scripts (mathematical analyses) or in Prism software (GraphPad; all other analyses). The Python scripts utilised several Python libraries: SciPy for statistical analysis (Virtanen et al., 2020), NumPy for numerical computations (Harris et al., 2020) and Matplotlib (Hunter, 2007) and Seaborn (Waskom, 2021) for visualisation. Statistical tests are defined in the figure legends and statistical significance was considered as *p* < 0.05. Figures were assembled in Adobe Illustrator.

### Code availability

The code used in this study is available on Github (https://github.com/ecuracosta/dorsal-ventral_gene_expression_in_the_regenerating_axolotl_spinal_cord) and Zenodo (Cura Costa and Chara, 2024).

## Results and discussion

### Mathematical modelling of dorsal-ventral domains during spinal cord regeneration

First, we confirmed the protein expression domains of SOX2 (expressed in neural progenitor cells) and the dorsal-ventral genes PAX7, PAX6 and SHH in steady state axolotl spinal cords (Figure 1a). Next, to profile the expression of these genes during spinal cord regeneration, we performed Hybridisation Chain Reaction (HCR) smFISH on tail sections harvested at steady state or at 14 days post-tail amputation (14 dpa) (Figures 1b-c). We included the roof plate gene *Msx1* in these assays as well as *Nkx6.1*, which had not been assayed previously in axolotls. We found *Nkx6.1* to be expressed in floor plate and ventro-lateral progenitors in a similar manner to the neural tube of mouse and chick (Figure 1c) (Briscoe et al., 2000; Qiu et al., 1998; Sander et al., 2000). Between these genes, we could identify at least 4 molecular domains whose arrangement appeared superficially similar between steady state and regeneration (from dorsal to ventral: *Msx1+Pax7+; Pax7+Pax6+; Pax6+Nkx6.1+; Nkx6.1+Shh+*) (Figure 1c).

**Figure 1.**
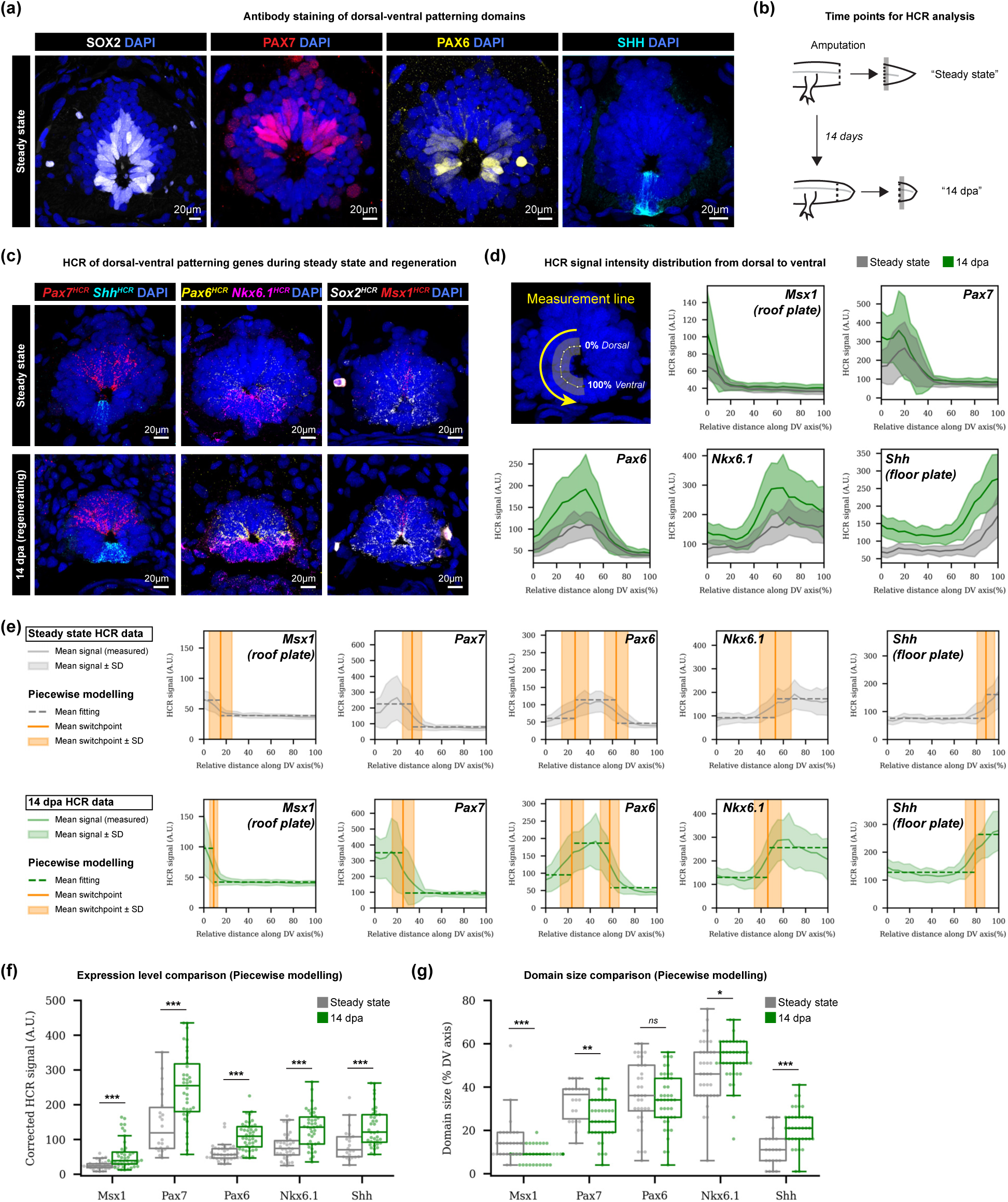
Mathematical modelling of dorsal-ventral gene expression during axolotl spinal cord regeneration. (a) Cross sections of steady state axolotl spinal cord, immunostained for neural progenitor gene (SOX2), dorsal-ventral transcription factors (PAX7, PAX6) or floor plate signal (SHH). DAPI labels nuclei. Maximum intensity projections through 20 mm of tissue, acquired with confocal microscopy. Dorsal is up and ventral is down. (b) Harvesting of steady state and 14 dpa regenerating spinal cord. Gray box indicates approximate analysis area. (c) Cross sections of spinal cords at steady state (top row) or 14 dpa (bottom row), stained using HCR for mRNA encoding dorsal-ventral patterning genes (*Msx1*, *Pax7, Pax6, Nkx6.1, Shh*) or neural progenitor gene *Sox2*. DAPI labels nuclei. Maximum intensity projections through 20mm of tissue, acquired with confocal microscopy. (d) Fluorescence intensity plots for HCR data at steady state and 14 dpa. *x*-axis is normalised distance along the dorsal-ventral (DV) axis, from dorsal to ventral. *y*-axis is HCR signal intensity (measured gray values), which was measured by using the segmented line tool (Fiji) to draw a line of thickness 100 through the neural progenitor layer of the spinal cord and using the Measurement function. *n* numbers are given in Table 1. (e) Plots depicting the fits of the piecewise models to the HCR data at steady state and 14 dpa. Solid lines and ribbons indicate mean HCR fluorescence measurements. Dotted lines indicate the mean fit of the piecewise model. Orange line indicates the switch point ± SD. (f) Box plots comparing the expression levels of dorsal-ventral genes at steady state and 14 dpa, as determined by piecewise modelling. Dots indicate values from individually fitted replicates. “Corrected HCR signal” is HCR signal intensity minus background intensity. ***: *p* < 0.05, Mann-Whitney U tests. Exact *p* values: *Msx1* (1.35 x 10^-4^), *Pax7* (3.40 x 10^-4^), *Pax6* (3.44 x 10^-8^), *Nkx6.1* (6.84 x 10^-5^), *Shh* (4.31 x 10^-4^). (g) Box plots comparing dorsal-ventral domain sizes at the lumen at steady state and 14 dpa, as determined by piecewise modelling. Dots indicate values from individually fitted replicates. Statistical comparison was performed by Mann-Whitney U tests. Exact *p* values: *Msx1* (2.08 x 10^-4^), *Pax7* (3.07 x 10^-3^), *Pax6* (ns), *Nkx6.1* (2.77 x 10^-2^), *Shh* (1.76 x 10^-4^).

**Table 1.** Samples quantified for mathematical modelling of HCR *in situ* staining. All quantifications were performed on maximum intensity projections of 20 μm thick spinal cord cross-sections. *n* = 6 axolotls for each of steady state and 14 dpa.

| Gene | Condition | Number of sections quantified |
| --- | --- | --- |
| *Msx1* | Steady state | 30 |
|  | 14 dpa blastema | 36 |
| *Pax7* | Steady state | 22 |
|  | 14 dpa blastema | 38 |
| *Pax6* | Steady state | 36 |
|  | 14 dpa blastema | 43 |
| *Nkx6.1* | Steady state | 36 |
|  | 14 dpa blastema | 43 |
| *Shh* | Steady state | 22 |
|  | 14 dpa blastema | 38 |

How the expression levels and domain sizes of the dorsal-ventral genes change during axolotl spinal cord regeneration is not known. To gain insights into these processes, we measured HCR signal along a continuous dorsal-to-ventral line drawn through progenitors contacting the spinal cord lumen (Figure 1d) (number of quantified sections is indicated in Table 1). We then used mathematical modelling to quantify HCR signal profiles and compare the two conditions (steady state and regeneration) in an unbiased manner. Previously, we analysed cell cycle dynamics in the regenerating axolotl spinal cord using a piecewise model, which assumes that zones of homogeneous behaviour are separated by sharp boundaries (switch points) (Cura Costa et al., 2021; Rost et al., 2016). We reasoned that piecewise modelling could similarly extract ‘gene expression level’ and ‘domain size’ from the HCR data, with an attractive feature being that the switch point unambiguously determines the gene expression boundary for further analyses (Figure S1a). We modelled *Msx1, Pax7, Nkx6.1* and *Shh* using a 2-zone model, with the 2 zones representing ‘expression’ or ‘background’. For *Pax6*, whose expression occurs centrally in the dorsal-ventral axis, we used a 3-zone model (‘background’-‘expression’-‘background’) with two switch points corresponding to the dorsal and ventral limits of *Pax6* expression. We performed individual fitting of replicates (Supplementary Information), generated mean fittings (Figure 1e, Figure S1b) and then used these to derive values for gene expression level and domain size (Table 2).

**Table 2.** Mean HCR signal and domain sizes calculated by the piecewise model. Mean HCR signal is after correction by subtracting background fluorescence.

| Gene | Condition | Piecewise model |  |
| --- | --- | --- | --- |
| | | Mean HCR signal* ± SD | Mean domain size ± SD |
| *Msx1* | Steady state | 25.5 ± 12.4 | 15.2 ± 10.3 |
| | 14 dpa | 55.5 ± 41.0 | 9.0 ± 3.6 |
| *Pax7* | Steady state | 146.0 ± 88.7 | 33.5 ± 8.9 |
|  | 14 dpa | 255.3 ± 117.5 | 25.4 ± 9.9 |
| *Pax6* | Steady state | 60.4 ± 24.8 | 36.9 ± 15.3 |
|  | 14 dpa | 109.8 ± 39.9 | 34.0 ± 12.6 |
| *Nkx6.1* | Steady state | 79.8 ± 37.0 | 47.1 ± 14.3 |
|  | 14 dpa | 126.5 ± 61.1 | 53.9 ± 12.2 |
| *Shh* | Steady state | 85.6 ± 46.8 | 11.5 ± 8.2 |
|  | 14 dpa | 135.8 ± 6.3 | 21.1 ± 8.8 |

Piecewise modelling revealed a significant increase in the expression of all genes assayed from steady state to regeneration, with a mean HCR signal increase ranging from 1.6-fold (*Shh, Nkx6.1*) to 2.2-fold (*Msx1*) (Figure 1f). The calculated switch points revealed that the representation of dorsal-ventral progenitors contacting the spinal cord lumen changed during regeneration. The dorsal gene domains became smaller (*Msx1*: -40.7%*, Pax7*: - 24.1%), the lateral *Pax6* domain remained the same size and the ventral gene domains became larger (*Nkx6.1*: +14.4% and *Shh*: +84.5%) at 14 dpa compared to steady state (Figure 1g). Thus, this analysis suggested that the roof plate and floor plate signalling centres were the domains that changed the most in their representation at the lumen. We confirmed the increase in *Shh*+ floor plate size through an independent assay, immunostaining for SHH protein at steady state and at 14 dpa (Figures S1c-d).

In summary, through mathematical modelling of HCR data, we determined that axolotl spinal cord regeneration proceeds *via* significant increases in the expression level of dorsal-ventral genes and a re-distribution of progenitor domains at the spinal cord lumen. Notably, we found that the floor plate signalling centre increases in size during axolotl spinal cord regeneration.

### Live labelling reveals changes in floor plate dynamics in the anterior-posterior axis

How do these dorsal-ventral changes relate to the anterior-posterior (snout-to-tail) axis of the regenerating spinal cord, particularly in the region of injury-activated progenitor cells (Albors et al., 2015; Cura Costa et al., 2021; Rost et al., 2016)? With the aim of resolving such dynamics live during regeneration, we designed a dual reporter axolotl to co-visualise dorsal cells and floor plate in which *Pax7* and *Shh* regulatory sequences controlled the expression of mCherry and EGFP fluorescent proteins respectively.

We previously generated a *Pax7* knock-in axolotl that co-expresses membrane-targeted mCherry and tamoxifen-inducible Cre recombinase from the *Pax7* locus (“*Pax7^mCherry-dERCre^*”, (Fei et al., 2017)). We used the same strategy to generate a *Shh* knock-in axolotl that expresses membrane-targeted EGFP and tamoxifen-inducible Cre from the *Shh* locus (“*Shh^EGFP-dERCre^*”). We mated these axolotls together to generate germline-transmitted dual reporter axolotls that simultaneously label *Pax7*+ and *Shh*+ cells with distinct fluorophores (Figure 2a, top). Transgenic axolotls used in this work are listed in Table 3.

**Figure 2.**
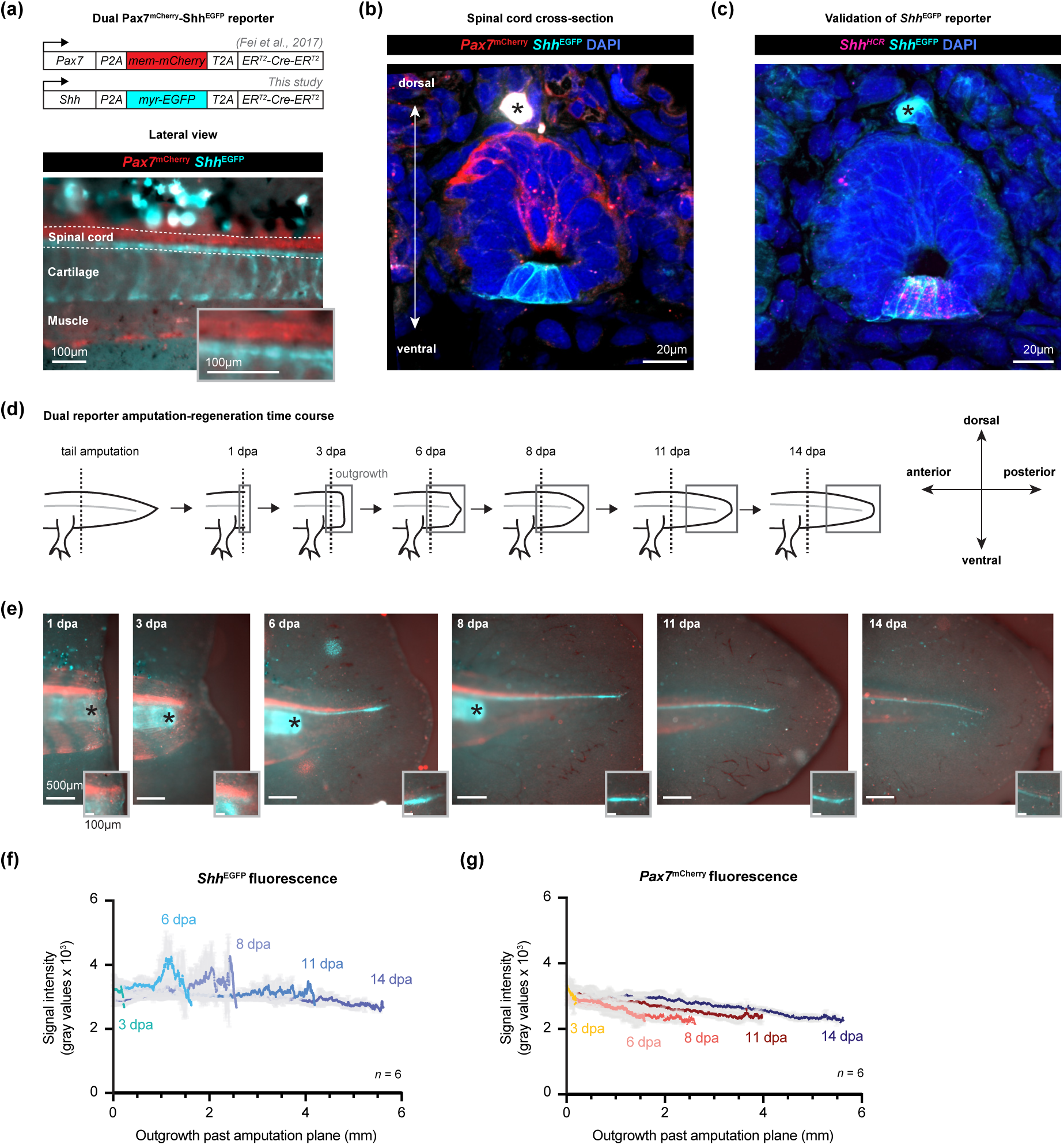
Live tracking of a dorsal-ventral reporter axolotl reveals a high *Shh* upregulation zone during regeneration. (a) A dual transgenic axolotl to track dorsal-ventral gene expression in axolotl spinal cord. CRISPR/Cas9-mediated knock-in results in co-expression of mCherry and tamoxifen-inducible Cre from the *Pax7* locus (Fei et al., 2017) or EGFP and tamoxifen-inducible Cre from the *Shh* locus (this study). Dual transgenics are heterozygous for each allele. A lateral view of a 2 cm axolotl reveals mCherry and EGFP expression on the dorsal and ventral sides of the spinal cord respectively. Additional expression is seen in muscle cell lineages (mCherry) and the cartilage rod (EGFP). (b) Spinal cord cross section from a 5 cm dual transgenic axolotl at 14 dpa. Red and cyan depict endogenous *Pax7^mCherry-dERCre^* and *Shh^EGFP-dERCre^* fluorescence. DAPI labels nuclei. Asterisk indicates autofluorescence. Maximum intensity projection through 20mm of tissue, acquired with confocal microscopy. (c) Spinal cord cross section from a 5 cm dual transgenic axolotl at 14 dpa. *Pax7^mCherry-dERCre^*is not depicted. Cyan depicts endogenous *Shh^EGFP-dERCre^* fluorescence. Magenta is HCR labelling against endogenous *Shh* mRNA. 100% of *Shh^EGFP-dERCre^+* cells were *Shh* mRNA+ (*n* = 72 cells from 6 spinal cords). Asterisk indicates autofluorescence. Maximum intensity projection through 20 mm of tissue, acquired with confocal microscopy. (d) An amputation-regeneration time course to measure changes in *Pax7^mCherry-dERCre^* and *Shh^EGFP-dERCre^* expression. Boxed areas represent images areas in (e). (e) Widefield microscopy of regenerating tails from 3 cm axolotls. Insets are magnifications of the regenerating spinal cord tip. Asterisks indicate the amputated tip of the cartilage rod, which acts as an indicator of the amputation plane. (f) Quantification of *Shh^EGFP-dERCre^* fluorescent signal in the regenerating part of the spinal cord. Dark lines are mean intensity values averaged from *n* = 6 spinal cords per time point; pale lines indicate standard deviations. (g) Quantification of *Pax7^mCherry-dERCre^* fluorescent signal in the regenerating part of the spinal cord. Dark lines are mean intensity values averaged from *n* = 6 spinal cords per time point; pale lines indicate standard deviations.

**Table 3.**
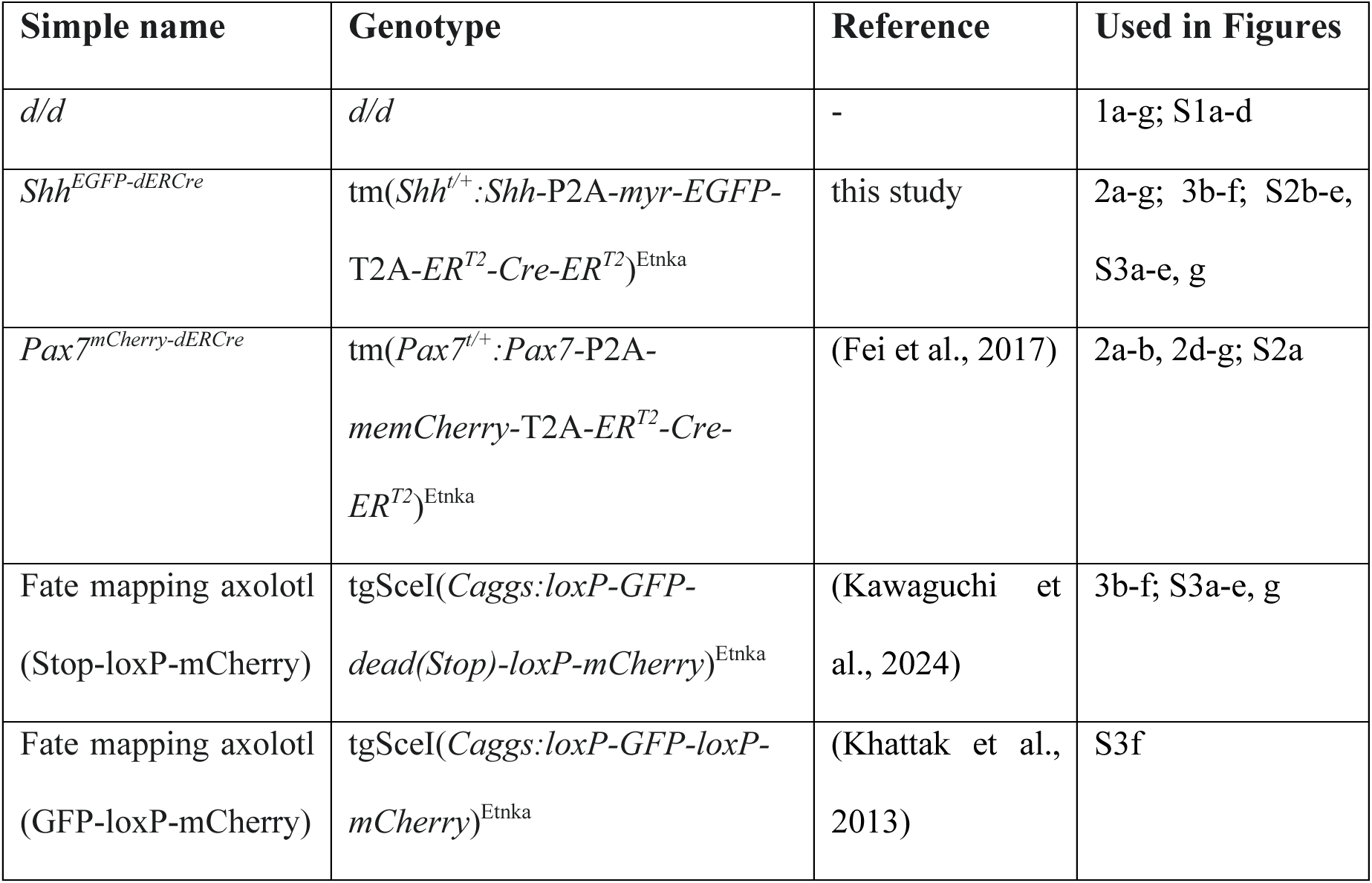
Transgenic axolotls used in this study.

Imaging dual reporter axolotls revealed restriction of mCherry and EGFP to opposite sides of the spinal cord, suggesting correct labelling of dorsal and ventral progenitors (Figure 2a, bottom and Figure 2b). We demonstrated previously, and confirm here (Figure S2a), that *Pax7^mCherry-dERCre^* faithfully recapitulates *Pax7* expression in the spinal cord and in tail muscle (Fei et al., 2017). We similarly tested the fidelity of the *Shh* reporter by performing HCR against *Shh* transcripts on *Shh^EGFP-dERCre^* spinal cord cross sections (Figure 2c). We found that 100% of EGFP+ cells expressed *Shh* transcripts (*n* = 72 cells from 6 spinal cords), although not all *Shh*+ cells expressed EGFP (67.5 ± 14.0 % of *Shh*+ cells were EGFP+, Figure S2b). The mean EGFP expression level was 2.1 times in regenerating tails compared to steady state (Figures S2c-d), a magnitude consistent with our *Shh* HCR analyses (Figure 1f).

Having validated the dual reporter axolotls, we performed tail amputation and live imaged the regenerating spinal cord every 2-3 days until 14 dpa (Figure 2d). We observed fluorescence in the outgrowing spinal cord and, consistent with the fixed tissue data, mCherry and EGFP appeared restricted to the dorsal and ventral sides (Figure 2e). Interestingly, this time series revealed a transient and spatially restricted increase in *Shh^EGFP-dERCre^* signal towards the spinal cord tip (also called ‘terminal vesicle’) (Figure 2e, insets). This high signal zone was located more posteriorly in the spinal cord than the regions harvested for the HCR analyses. Thus, we infer that, in addition to a general increase in *Shh* expression during regeneration (Figure 1f), there is a posterior zone in which *Shh*^EGFP^ signal is particularly high. This high signal could be the result of elevated *Shh* expression, a higher density of *Shh*+ cells, or a combination of both. By measuring mean fluorescence intensity in outgrowing spinal cords (Figure S2e), we found that this high *Shh^EGFP^* signal zone extended ∼800 μm anteriorly from the regenerating tip and was apparent at 6-8 dpa, before disappearing by day 14 (Figure 2f). An equivalent analysis of *Pax7^mCherry-dERCre^* revealed no such dynamics – and, in fact, there was a tendency of decreasing expression towards the tail tip across all time points (Figure 2g). In sum, we identified both anterior-posterior and dorsal-ventral changes in floor plate dynamics during axolotl spinal cord regeneration.

### *Shh*+ cells selectively generate *Shh*+ cells during spinal cord regeneration

Given these spatiotemporal differences in floor plate behaviour, an important question is how *Shh*+ cells contribute to the regenerating spinal cord. A simple model is that *Shh*+ cells give rise only to *Shh+* floor plate during regeneration. However, another possibility is that *Shh*+ floor plate can switch dorsal-ventral identity to give rise to other neural progenitors (Mchedlishvili et al., 2007). To distinguish between these possibilities (fate-restricted model vs. flexible identity model, Figure 3a), we used a genetic strategy to label *Shh*+ cells and track their progeny during regeneration. We crossed *Shh^EGFP-dERCre^* axolotls, which express tamoxifen-inducible Cre recombinase, to our previously published fate mapping axolotl (*Caggs*:*loxP-Stop-loxP-mCherry*) (Figure 3b). Treating the progeny with 4-hydroxytamoxifen (4-OHT) induces recombination and removal of the Stop cassette, labelling *Shh*+ cells and their progeny permanently with mCherry.

**Figure 3.**
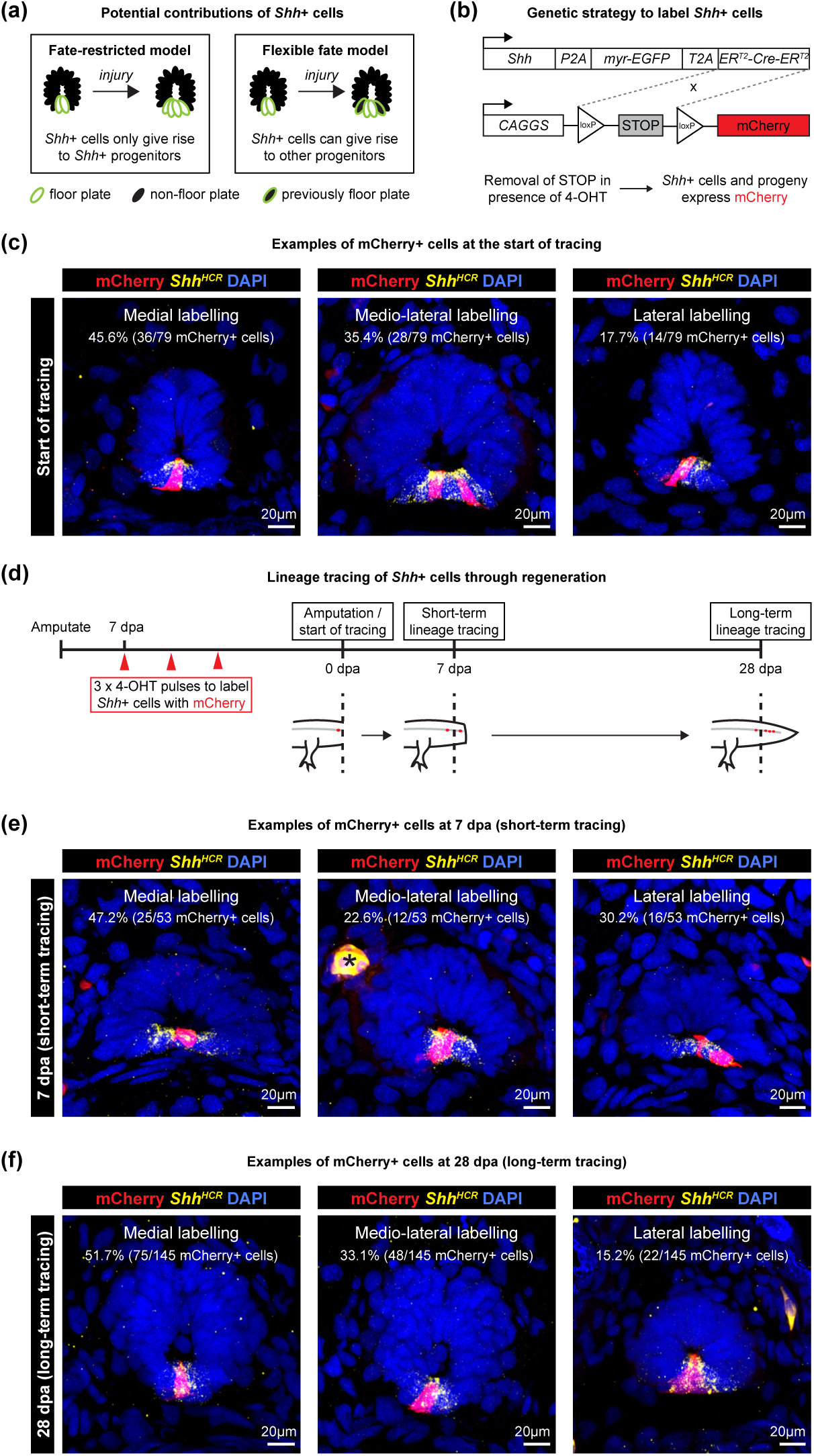
*Shh+* cells give rise to *Shh+* cells during spinal cord regeneration. (a) Two hypotheses for how *Shh*+ cells contribute to spinal cord regeneration. (b) Genetic strategy to lineage trace *Shh+* cells. *Shh*+ cells continuously express tamoxifen-inducible Cre. Pulse application of 4-OHT induces Cre translocation to the nucleus, where it excises the STOP sequence in the fate mapping cassette by Cre/loxP recombination. This results in permanent expression of mCherry in the *Shh*+ cell and its progeny. (c) Spinal cord cross sections from *Shh* lineage tracing axolotls pulsed three times with 4-OHT. 98.7% of mCherry cells express *Shh* mRNA (*n* = 78/79 cells, from 15 tails). Cells were labelled in all regions (medial, medio-lateral, lateral) of the floor plate. DAPI labels nuclei. Maximum intensity projections through 20 mm of tissue, acquired with confocal microscopy. (d) Amputation-regeneration time course to lineage trace *Shh*+ cells. *Shh*+ cells were labelled with mCherry as in (c), then spinal cords were re-amputated within 500 mm of labelled cells to induce them to contribute to regeneration. Replicate spinal cords were harvested at 7 dpa (short-term tracing) and 28 dpa (long-term tracing) to assess lineage contributions. (e) Spinal cord cross sections harvested from *Shh* lineage tracing axolotls at 7 dpa (short-term tracing). 100% of mCherry cells expressed *Shh* mRNA (*n* = 53 cells from 19 spinal cords). Labelled cells were seen in all regions of the floor plate. DAPI labels nuclei. Asterisk indicates autofluorescence. Maximum intensity projections through 20 mm of tissue, acquired with confocal microscopy. (f) Spinal cord cross sections harvested from *Shh* lineage tracing axolotls at 28 dpa (long-term tracing). 100% of mCherry cells express *Shh* mRNA (*n* = 145 cells from 8 spinal cords). Labelled cells were seen in all regions of the floor plate. DAPI labels nuclei. Maximum intensity projections through 20 mm of tissue, acquired with confocal microscopy.

Initially, we attempted lineage labelling at steady state by treating axolotls once or three times with 2 μM 4-OHT, but neither strategy induced mCherry labelling robustly (Figure S3a), potentially due to low expression of *Shh* and *Cre* at steady state (Figures S2c-d). Therefore, we treated animals with 4-OHT after tail amputation, which elevates *Shh* expression (Figures S2c-d). By treating animals three times with 4-OHT from 7 dpa, we succeeded in labelling sparse ventral cells in the spinal cord (“start of lineage tracing”) (Figure S3a). Importantly, this labelling only occurred in 4-OHT treated animals (Figure S3b). HCR for *Shh* mRNA revealed that almost 100% of mCherry-labelled cells were *Shh*+ (*n* = 78/79 cells from 15 tails) (Figure 3c). Notably, we labelled medial, medio-lateral and lateral floor plate cells, allowing us to trace all regions of the floor plate (Figure 3c). Across all samples, we observed only one single mCherry+ *Shh-*negative cell, demonstrating the overall specificity of labelling (Figure S3c).

Having labelled *Shh+* cells, we examined their lineage contributions to spinal cord regeneration. We re-amputated labelled tails within a zone 500 μm posterior to mCherry+ cells (Figure S3d), as neural progenitors within this zone contribute to spinal cord regeneration (Mchedlishvili et al., 2007). We harvested tails at 7 dpa (“short-term tracing”) or 28 dpa (“long-term tracing”) (Figure 3d). As expected, we observed an increase in the number of mCherry+ cells during the tracing window as they proliferated and contributed to regeneration (Figure S3e). To identify the traced cells, we performed HCR against *Shh*. We found that 100% of mCherry labelled cells expressed *Shh* mRNA both at 7 dpa (Figure 3e) (*n* = 53 cells from 19 tails) and at 28 dpa (Figure 3f) (*n* = 145 cells from 8 tails), with little change in the positions of the labelled cells within the floor plate (medial, medio-lateral or lateral) (Figures 3c, e-f). These results support that *Shh*+ cells maintain floor plate identity during axolotl spinal cord regeneration.

One risk with the lineage tracing was that we failed to label other progenitor cells due to a lack of expression of the fate mapping cassette. To exclude this possibility, we analysed spinal cord sections from *Caggs:loxP-GFP-loxP-mCherry* fate mapping axolotls (Khattak et al., 2013). These axolotls use the same expression system as those used in our lineage tracings but additionally express GFP in any cell that expresses the memory cassette (independent of Cre/loxP recombination). We found that whenever a *Shh*+ cell expressed the memory cassette, neighbouring (more dorsal) progenitors also expressed the memory cassette, indicating the potential to become labelled (*n* = 27/28 sections analysed, harvested from 6 axolotls) (Figure S3f). On the other hand, we found that the most dorsal progenitors on the other side of the spinal cord frequently lacked expression of the fate mapping cassette (Figure S3f). This is an important consideration for investigations into dorsal cells.

As a result of the sparse labelling efficiency in these experiments, we could detect that *Shh*+ cells change morphology during regeneration. At the 28 dpa time point, anterior spinal cord regions (closer to the original amputation plane) had already regenerated neurons while posterior regions (towards the outgrowing tip) still lacked neurons (Figure S3g). As neurons are regenerated in an anterior-to-posterior direction, the anterior regions containing neurons could be considered more ‘mature’ regenerate tissue compared to the more ‘immature’ posterior regions lacking neurons. We found that *Shh*+ cells in the immature spinal cord had a simple, trapezoid morphology, while *Shh*+ cells in the mature part had a more complex shape including an apical process extending towards the spinal cord lumen and one or more basal processes ventrally (Figure S3g). This morphological difference is likely related to maturation state rather than anterior-posterior differences, as all labelled *Shh*+ cells had the simpler morphology at the 7 dpa time point (Figure 3e). This is the first time that the complex morphology of floor plate cells has been captured in regenerating spinal cord.

Several injury paradigms are used to study mechanisms of spinal cord regeneration. Spinal cord transection in zebrafish elevates dorsal-ventral patterning gene expression, including *Shh,* local to the injury site (Reimer et al., 2009). In this study, by taking a mathematical modelling approach to the axolotl model, we found that tail amputation triggers not only a general increase in dorsal-ventral gene expression but also a larger floor plate and a high *Shh^EGFP^* zone within ∼800 μm of the regenerating spinal cord tip. The function of this high *Shh^EGFP^* zone, and if it relates to previous suggestions of a higher plasticity in progenitor identity at the terminal vesicle (Mchedlishvili et al., 2007) are important topics for future study. In both zebrafish and axolotls, upregulation of dorsal-ventral genes is observed by 14 days post-injury. It is possible that this upregulation reflects the acquisition of a more development-like cellular state for regeneration. Interestingly, *Pax6* is upregulated after spinal cord transection in rats concomitant with cell proliferation (Yamamoto et al., 2001), suggesting the potential for similar (but more limited) molecular changes in mammals.

Although *Shh*+ cells persist in the axolotl spinal cord throughout life, their cellular contributions to regeneration had not been identified. By performing genetic lineage tracing, we found that *Shh*+ cells are limited to generating regeneration floor plate in the tail amputation model. We were only able to label sparse *Shh*+ cells due to a poor efficiency of Cre/loxP-mediated memory cassette recombination. It is likely that labelling efficiency could be improved by increasing Cre activity (e.g. reducing the number of ER^T2^ domains fused to the Cre recombinase) and/or increasing Cre expression level (e.g. expressing the Cre recombinase prior to EGFP in the knock-in cassette). However, sparse labelling was powerful for revealing floor plate cell morphology. Although floor plate cells are commonly described to be cuboidal or trapezoid, they are thought to have a more complex morphology characterised by apical and basal cellular processes (Campbell and Peterson, 1993; Yaginuma et al., 1991). Recently, it was found that the basal processes of chick floor plate cells comprise multiple extensions that enwrap the growth cones of dorsal commissural neurons and constrain them to a straight trajectory path (Ducuing et al., 2020). Here, we discovered that axolotl floor plate cells lack complex basal processes at 7 dpa but elaborate these later during regeneration (by 28 dpa), possibly reflecting functional maturation. These basal processes could serve an axon guidance function for regenerating axons, but this should be tested functionally.

Previous experiments had suggested that neural progenitors can change dorsal-ventral identity during axolotl regeneration (Mchedlishvili et al., 2007). Determining if *Shh*+ cells are exceptions to this behaviour, or if their fate could be changed by external manipulation or a different injury paradigm, are important future directions. Reciprocally, lineage tracing the other neural progenitors in the spinal cord will reveal the degree to which these cells can contribute to the formation of floor plate and roof plate signalling centres. During axolotl limb regeneration, cells that did not previously express *Shh* can readily generate *Shh*+ signalling centre cells (Otsuki et al., 2023). Studying the origins and fate limitations of signalling centres in different tissues will uncover different avenues to tissue patterning *in vivo* and in tissue engineering applications.

## Author contributions

L.I.A. designed experiments, performed all experiments, analysed data and wrote the manuscript. E.C.C. analysed data and wrote the manuscript. O.C. secured funding, analysed data and wrote the manuscript. L.O. conceived the project, secured funding, designed experiments, generated transgenic axolotls, analysed data, supervised the project and wrote the manuscript. E.M.T. conceived the project, secured funding, supervised the project and wrote the manuscript. All authors approved the manuscript.

## Acknowledgements

We thank Tanaka laboratory members for reagents (Hannah Stuart, Wouter Masselink, Anastasia Polikarpova, Pietro Tardivo) and project discussions (Elad Bassat, Katharina Lust, Hannah Stuart). We are grateful to the caretaker team (Magdalena Blaschek, Emina Silic, Victoria Szilagyi, Andrea Lentz-Koblenc, Veronika Vojnicsek) for excellent axolotl care and the Molecular Biology Service and BioOptics facility at IMP/IMBA for expert support.

E.CC. and O.C. were funded by a PICT-2019-03828 grant from AGENCIA (National Agency from the Promotion of Research, Technological Development and Innovation, Argentina). L.O. was funded by fellowship LT000785/2019-L from HFSP (Human Frontier Science Program). L.I.A. and E.M.T. were funded by Advanced Grant 742046 (RegGeneMems) from the ERC (European Research Council) and Special Research Program of the Austrian Science Fund (FWF) Project F78. For the purpose of open access, the authors have applied a CC BY public copyright licence to any Author Accepted Manuscript version arising from this submission.

