## Supplementary Figures 1-3 for "Lineage tracing of *Shh+* floor plate cells and dynamics of dorsal-ventral gene expression in the regenerating axolotl spinal cord"

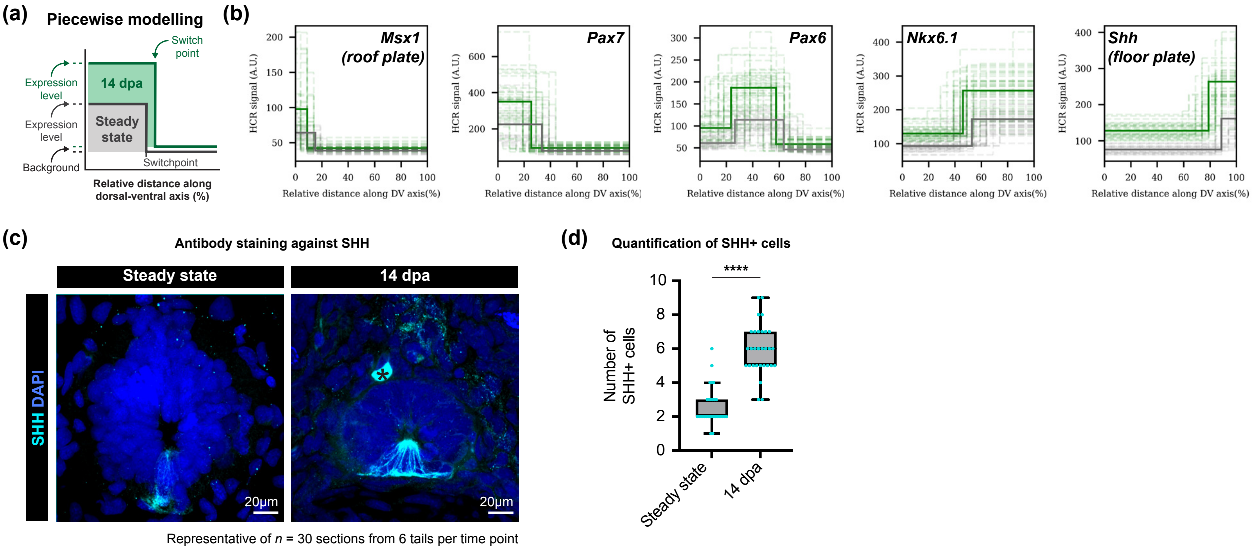


**Figure S1. Quantification of changes in dorsal-ventral patterning gene expression during regeneration.**
(a) Schematic depicting how the fit and switch point of piecewise modelling relate to gene expression level and domain size determination.
(b) Piecewise modelling of HCR fluorescence data. For each gene, the steady state fit is plotted with a solid gray line and 14 dpa fit with a solid green line. Individual fits to replicates are plotted with dotted lines.
(c) Spinal cord cross sections at steady state and 14 dpa, immunostained for SHH. DAPI labels nuclei. Asterisk indicates autofluorescence. Maximum intensity projections through 20 μm of tissue, acquired with confocal microscopy.
(d) Box plots showing the number of SHH+ cells at steady state and 14 dpa, as assessed by antibody staining. ****: $p = 1.79 \times 10^{-9}$, Kolmogorov-Smirnov test, n = 30 cross sections quantified per time point, harvested from 6 tails each.

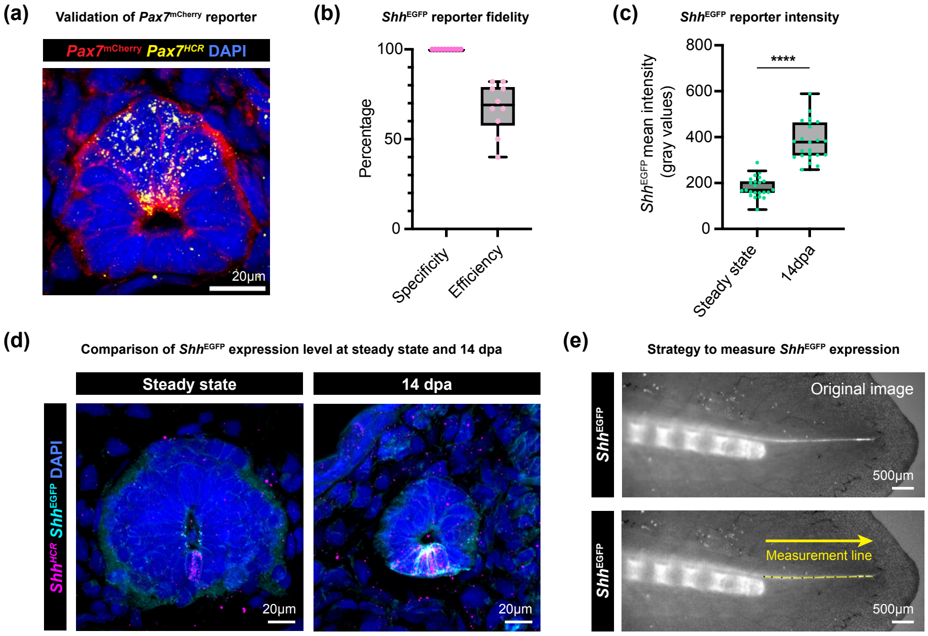


**Figure S2. Characterisation of Pax7$^{mCherry-dERCre}$-Shh$^{EGFP-dERCre}$ dual transgenic axolotl.**
(a) Spinal cord cross section from a 4 cm dual transgenic axolotl at 14 dpa. Shh$^{EGFP-dERCre}$ is not depicted. Red depicts endogenous Pax7$^{mCherry-dERCre}$ fluorescence. Yellow is HCR staining against Pax7 mRNA. Maximum intensity projection through 20 μm of tissue, acquired with confocal microscopy.
(b) Box plots depicting the fidelity of the Shh$^{EGFP-dERCre}$ reporter, as assessed in spinal cords at 14 dpa. Specificity: the percentage of Shh$^{EGFP-dERCre}$ cells that express Shh mRNA (assessed by HCR). Efficiency: the percentage of Shh mRNA-expressing cells that are positive for Shh$^{EGFP-dERCre}$. n = 10 spinal cords.
(c) Box plots depicting the signal intensity of the Shh$^{EGFP-dERCre}$ reporter in the floor plate at steady state vs 14 dpa. ****: $p = 1.58 \times 10^{-11}$, unpaired t-test with Welch's correction. n = 23 sections analysed per time point.
(d) Spinal cord cross sections from 4 cm dual transgenic axolotls at steady state (left) or at 14 dpa (right). Images are displayed with the same intensity settings. Pax7$^{mCherry-dERCre}$ is not depicted. Cyan is Shh$^{EGFP-dERCre}$ fluorescence. Magenta is HCR labelling against Shh mRNA (floor plate). Shh$^{EGFP-dERCre}$ expression is weaker at steady state than during regeneration. Maximum intensity projection through 20 μm of tissue, acquired with confocal microscopy.
(e) Strategy to quantify dual reporter fluorescence in the regenerating spinal cord. The segmented line tool (Fiji) was used to draw a line of thickness 10 through the outgrowing spinal cord from anterior to posterior, starting at the amputation plane (determined by the cartilage rod stump). The Measurement function was used to extract mCherry and EGFP fluorescence intensity as a continuous variable.

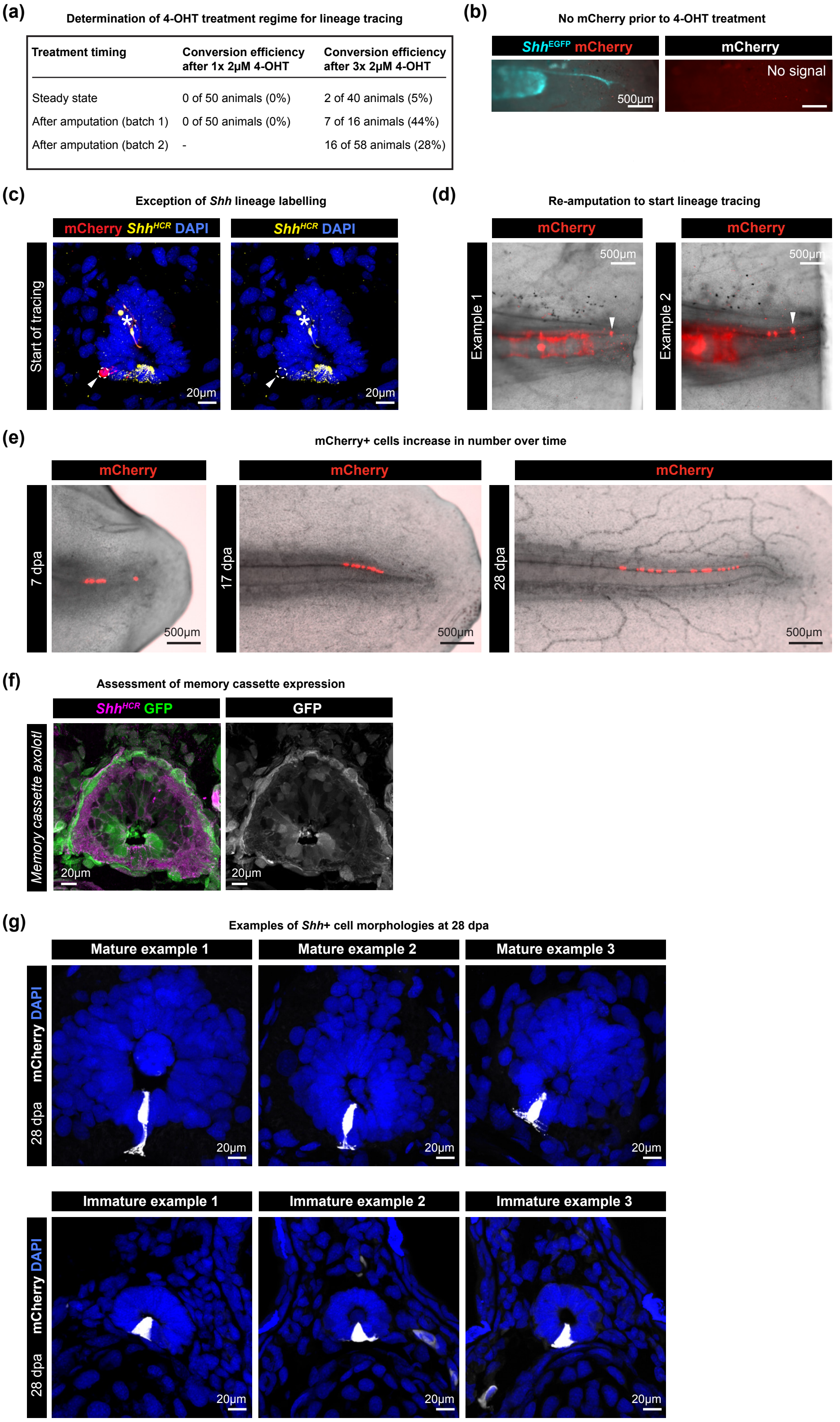


**Figure S3. Details of Shh lineage tracing experiment.**
(a) Induction of mCherry expression from the fate mapping cassette after different 4-OHT treatment conditions. Three overnight pulses of 4-OHT treatment from 7 dpa was the only condition that resulted in reliable mCherry labelling. In all experiments, approximately half of the animals would have inherited the memory cassette from the parent, resulting in a theoretical maximal labelling efficiency of ~50%.
(b) Lateral widefield image of a Shh lineage tracing axolotl prior to 4-OHT treatment. As expected, no mCherry labelling is present, while $Shh^{EGFP\text{-}dERCre}$ expression (cyan) can be seen in ventral cells.
(c) One single Shh negative cell labelled with mCherry (circled and arrowed) was observed in the “start of tracing” cohort (total 79 cells). Yellow is HCR against Shh mRNA. DAPI labels nuclei. Asterisk indicates autofluorescence.
(d) Lineage tracing was initiated by re-amputating 4-OHT-treated tails within a 500 μm zone posterior to mCherry-labelled cells (most posterior cell is arrowed in two examples).
(e) Lateral widefield images depicting expansion of mCherry-labelled cell clones in the spinal cord from 0 dpa to 28 dpa (long-term trace).
(f) Confirmation that the fate mapping cassette expresses in multiple domains of the spinal cord. Spinal cord cross section harvested from 3 cm Caggs:loxP-GFP-loxP-mCherry axolotls (Khattak et al., 2013) at 7 dpa. Green/gray is endogenous GFP (indicating fate mapping cassette expression). Whenever Shh+ cells (magenta) expressed the fate mapping cassette, adjacent spatial domains also expressed the cassette (n = 27 of 28 spinal cord sections). Maximum intensity projection through 20 μm of tissue, acquired with confocal microscopy.
(g) Examples of mCherry-labelled Shh+ cells at 28 dpa, located in the more mature part of the regenerate (indicated by the presence of peripheral neurons) or in the more immature part of the regenerate (indicated by lack of peripheral neurons and a small spinal cord diameter). Some Shh+ cells in the mature regenerate extended one or more protrusions from their ventral side, oriented laterally or directly away from the lumen (detected in 13/26 sections). Although Shh+ cells in the immature part of the regenerate also extended lateral processes (see Example 2), these were not as numerous as those in the mature part, and were not oriented away from the lumen (representative of 56 sections).
