## Supplementary Information for "Lineage tracing of *Shh+* floor plate cells and dynamics of dorsal-ventral gene expression in the regenerating axolotl spinal cord"

[1] Institute of Molecular Biotechnology of the Austrian Academy of Sciences (IMBA), Vienna BioCenter (VBC), Dr. Bohr-Gasse 3, 1030 Vienna, Austria
[2] Research Institute of Molecular Pathology (IMP), Vienna BioCenter (VBC), Campus-Vienna-Biocenter 1, 1030 Vienna, Austria
[3] Institute of Physics of Liquids and Biological Systems (IFLYSIB), National Scientific and Technical Research Council (CONICET), University of La Plata, La Plata B1900BTE, Argentina
[4] School of Biosciences, University of Nottingham, Sutton Bonington Campus, Nottingham LE12 5RD, United Kingdom
[5] Center for Information Services and High Performance Computing, Technische Universität Dresden, Dresden, Germany
[6] Instituto de Tecnología, Universidad Argentina de la Empresa, Buenos Aires, Argentina

### 1 Introduction

In this supplementary material, we provide a detailed explanation of the piecewise constant model used to analyse gene expression data along the dorsal-ventral (DV) axis of the spinal cord. Similar methodologies, including the use of step functions for data analysis, have been discussed in previous studies on spinal cord regeneration (Rost et al., 2016; Cura Costa et al., 2021). In addition, we describe the statistical analyses performed to compare gene expression levels and domain sizes between different conditions for each gene. The implementations of these analyses, using several libraries including NumPy for numerical calculations (Harris et al., 2020), Matplotlib (Hunter, 2007) and Seaborn (Waskom, 2021) for plotting, and SciPy for statistical analysis (Virtanen et al., 2020), can be found at the following link: GitHub Repository (Cura Costa and Chara, 2024).

### 2 Piecewise constant model fitting

We employed piecewise constant functions to model gene expression data along the DV axis (Figures 5.1). Two forms of the piecewise constant function were used: a two-step function for most genes and a three-step function specifically for Pax6.

#### 2.1 Two-step piecewise constant function

The two-step piecewise constant function is defined as follows:

$$f(x) = \begin{cases} a & \text{if } x < sp \\ b & \text{if } x \geq sp \end{cases} \tag{1}$$

where $a$ and $b$ are the HCR signal levels before and after the switchpoint (sp), respectively. The switchpoint is the relative position in the DV axis where the signal changes levels.

#### 2.2 Three-step piecewise constant function for Pax6

For Pax6, we used a three-step piecewise constant function, defined as follows:

$$f(x) = \begin{cases} a & \text{if } x < sp1 \\ b & \text{if } sp1 \leq x < sp2 \\ c & \text{if } x \geq sp2 \end{cases} \tag{2}$$

where $a$ is the constant level before the first switchpoint (sp1), $b$ is the constant level between the first switchpoint (sp1) and the second switchpoint (sp2), and $c$ is the constant level after the second switchpoint (sp2).

### 3 Sum of Squared Errors (SSE) analysis

To determine the optimal fit, we calculated the mean signal levels for the zones defined by the switchpoint in the two-step function (or pairs of switchpoints in the three-step function). We then computed the sum of squared errors (SSE) to evaluate the model's fit (Figures 5.2):

$$\text{SSE} = \sum_{i=1}^{n} (y_{\text{data},i} - f(x_{\text{data},i}, \text{params}))^2 \tag{3}$$

where $y_{\text{data}}$ represents the observed data points, $f(x_{\text{data}}, \text{params})$ represents the fitted model values, and $n$ is the number of data points.

### 4 Statistical analysis of gene expression data

To compare gene expression levels and domain sizes between different conditions, we derived data from the piecewise constant fitting models. The following steps outline the process:

#### 4.1 Data extraction for comparative analysis

For each gene and condition, the optimal switchpoints determined from the piecewise constant fitting were used to segment the data into distinct regions. The HCR signal levels in these regions were then averaged to obtain representative values for each segment. Specifically:

- For the two-step model, signal levels before and after the switchpoint were averaged.
- For the three-step model used for Pax6, signal levels were averaged for the three segments defined by the two switchpoints.

The resulting mean signal levels and the positions of the switchpoint(s) were used as inputs for comparative analysis.

#### 4.2 Statistical comparisons between conditions

To statistically compare gene expression levels and domain sizes between different conditions, we performed the following steps:

- Calculation of mean signal differences: for each replica, the differences in mean signal levels between expression and basal zones were calculated for each gen in both conditions (for Pax6, the mean of the two basal intensities was subtracted to correct the signal).
- Calculation of domain sizes: for each replica, the domain sizes were calculated as the two (or three) domains delimited by the switchpont(s) for each gen in both condtions.
- Statistical tests: we conducted statistical tests to evaluate the significance of the observed differences for each gen between conditions. The tests included a non-parametric test, the Mann-Whitney U test, and a parametric test, an independent t-test. This tests determines if there is a significant difference between the means of the two groups.

The results of these analyses were used to generate comparative figures (Figure 1f and Figure 1g in the main text), providing a visual representation of the differences in gene expression patterns between the studied conditions.

### 5 Figures

#### 5.1 Individual fittings

The observed data points in the figures represent the HCR signal levels along the dorsal-ventral axis. The best fit (orange dashed line) shows the signal levels between switchpoints. The switchpoint is the position where the signal level changes.

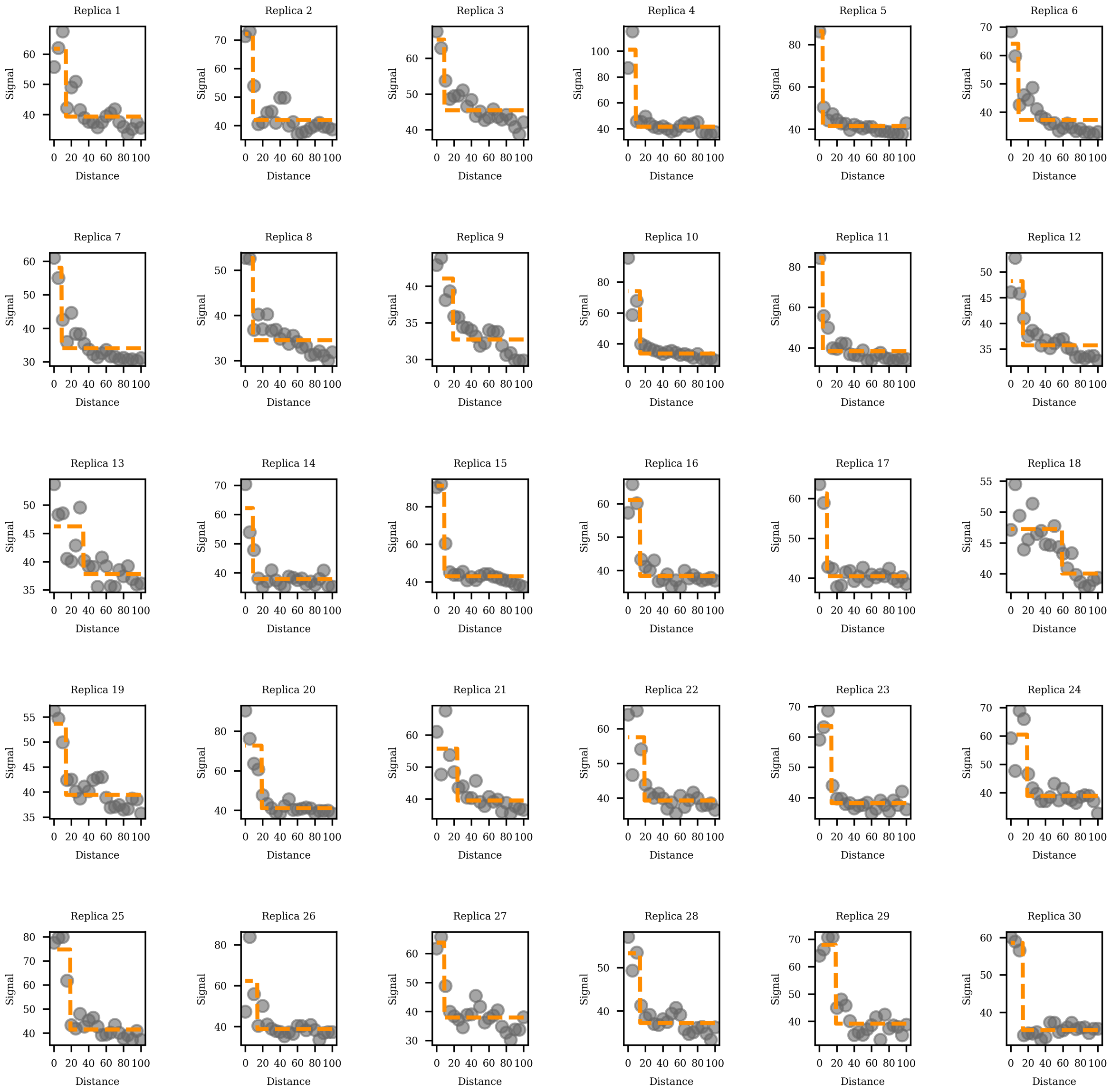

Figure 1: Individual fittings for Msx1, steady state.

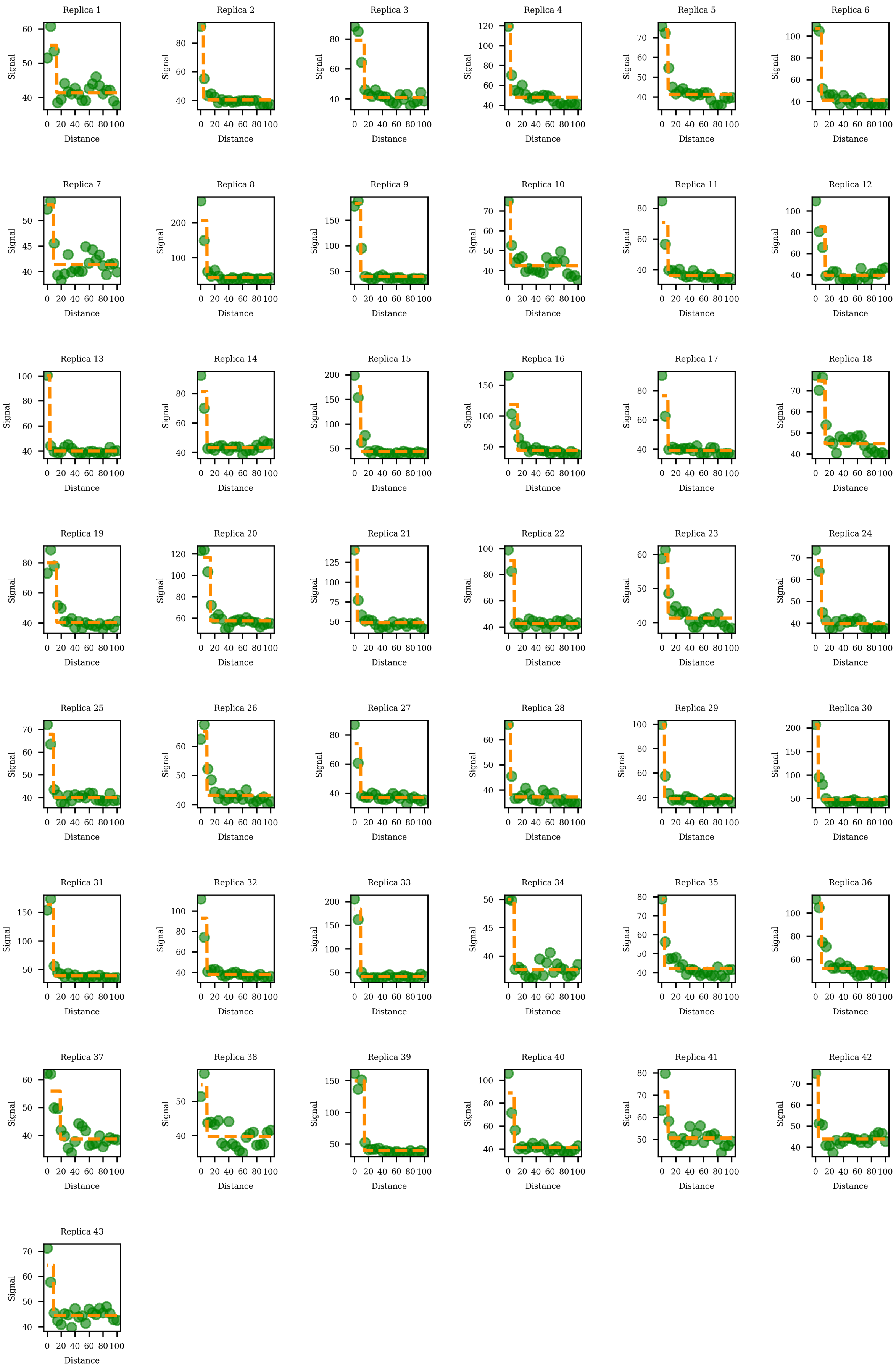

Figure 2: Individual fittings for Msx1, 14 dpa.

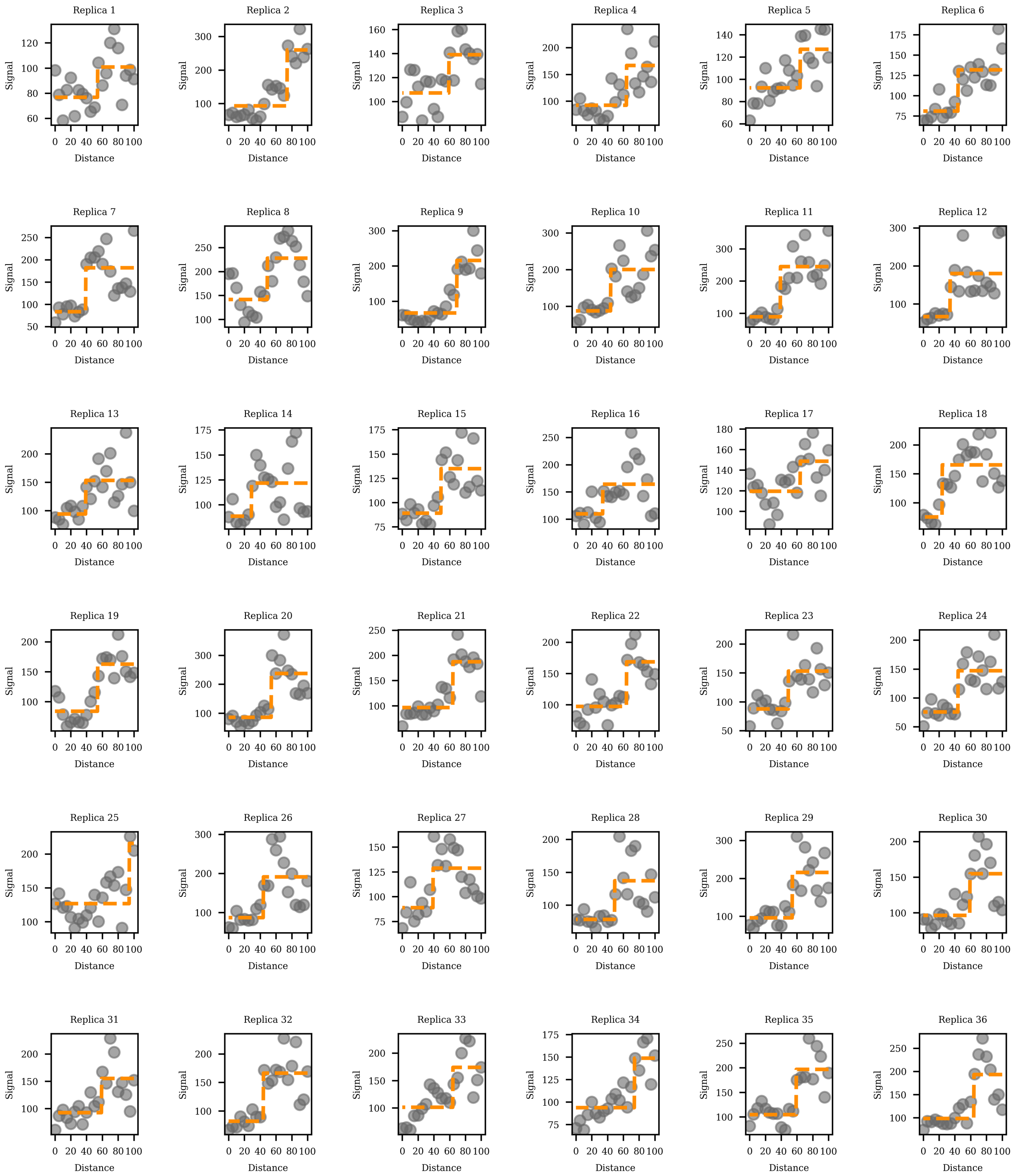

Figure 3: Individual fittings for Nkx6.1, steady state.

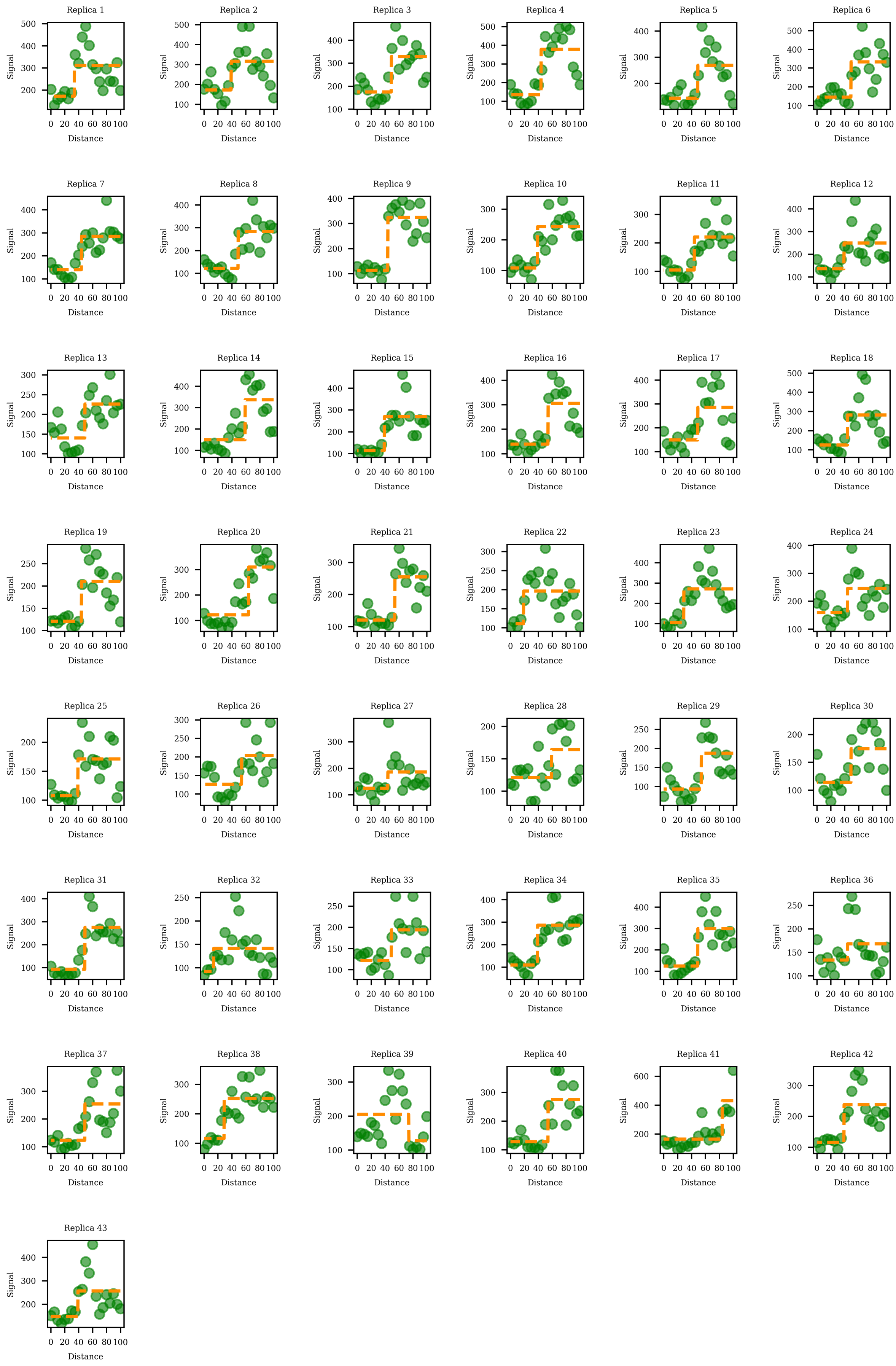

Figure 4: Individual fittings for Nkx6.1, 14 dpa.

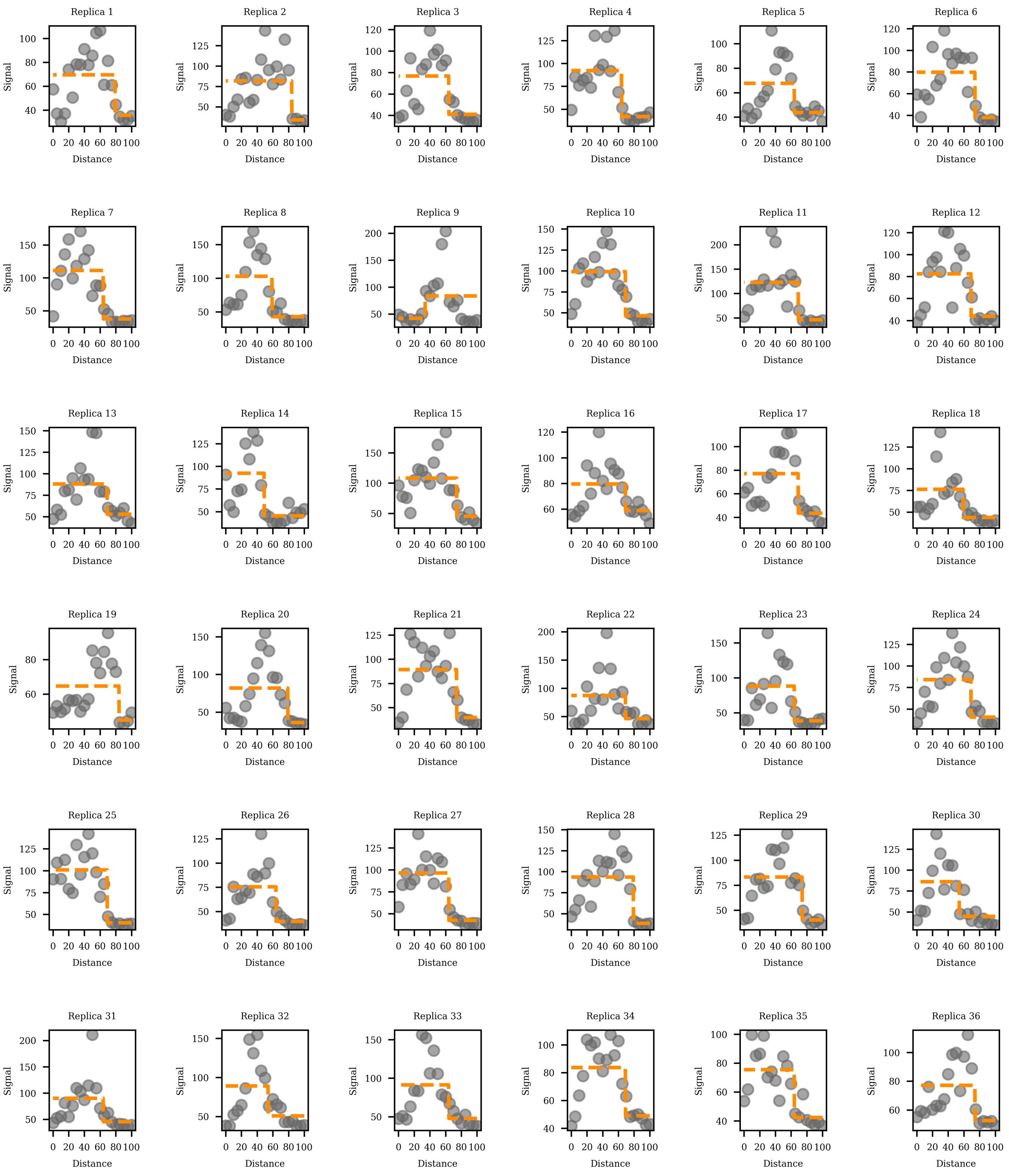

Figure 5: Individual fittings for Pax6, steady state.

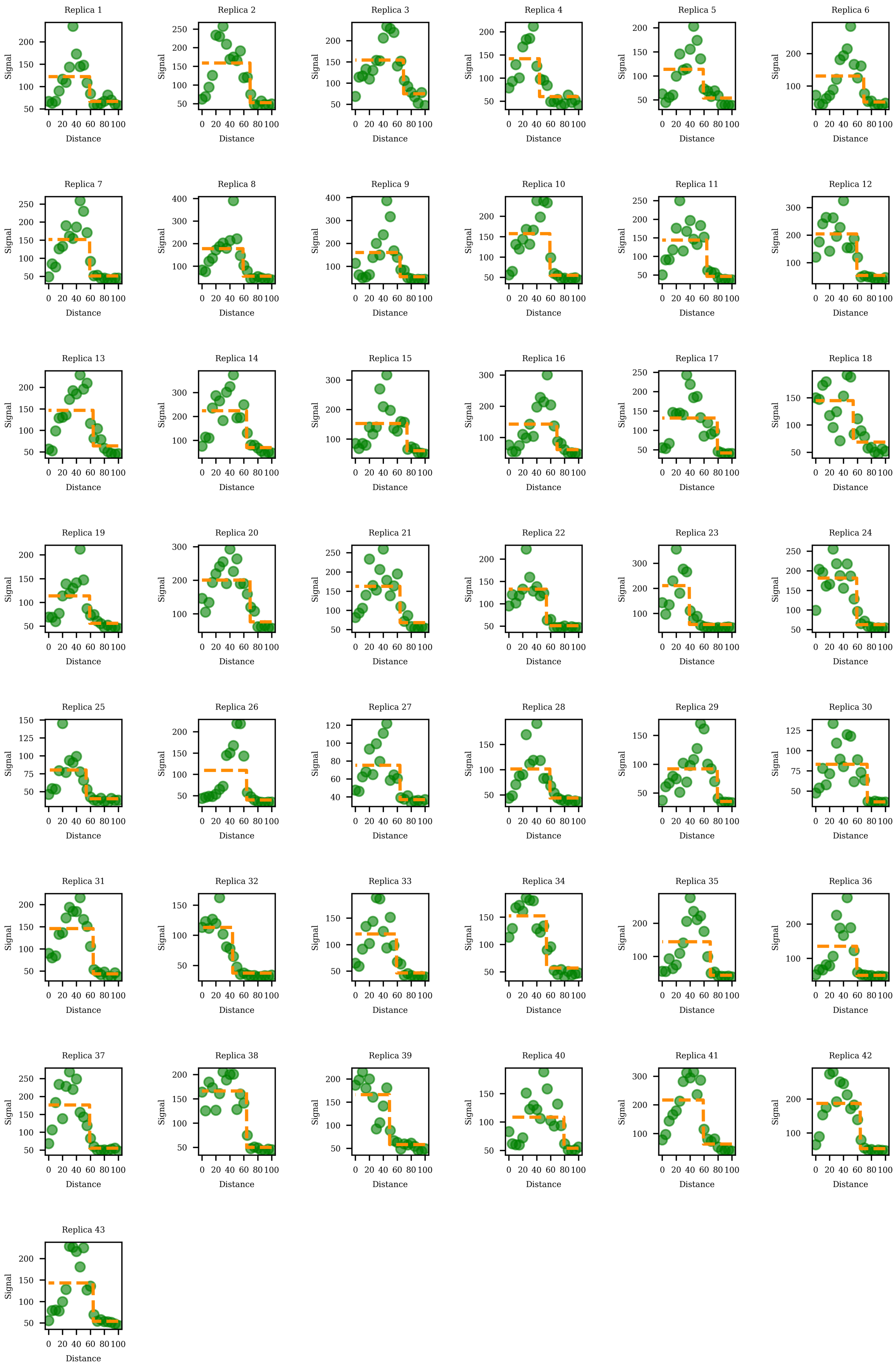

Figure 6: Individual fittings for Pax6, 14 dpa.

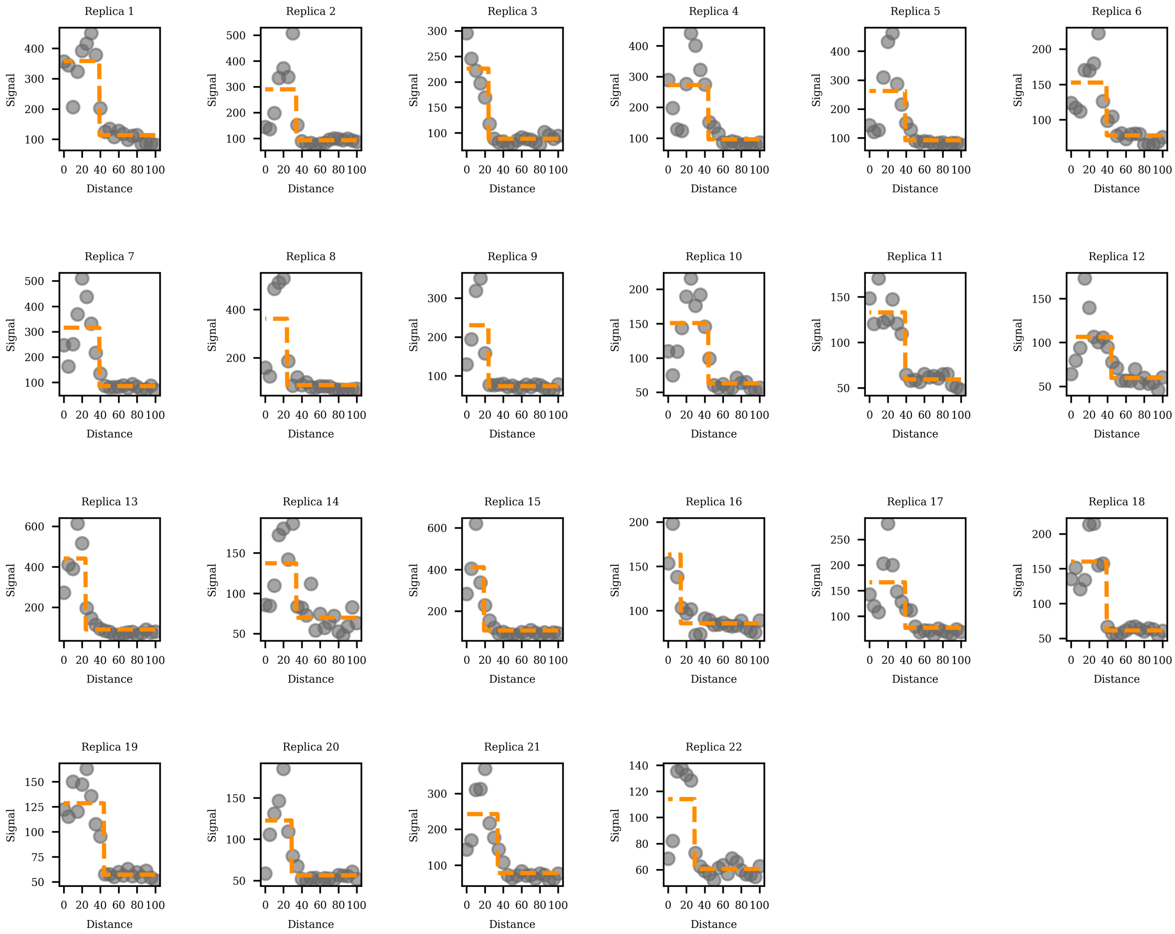

Figure 7: Individual fittings for Pax7, steady state.

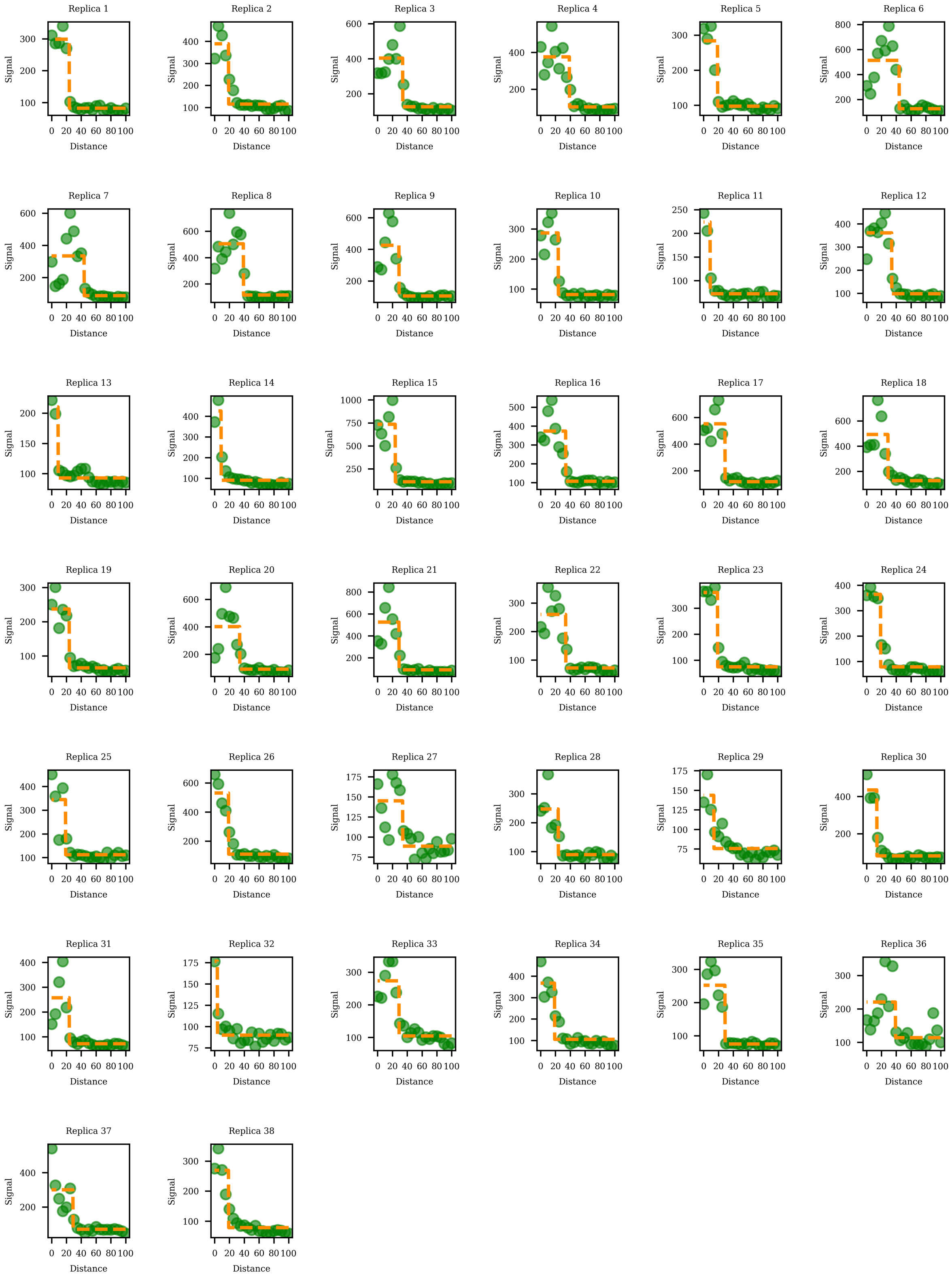

Figure 8: Individual fittings for Pax7, 14 dpa.

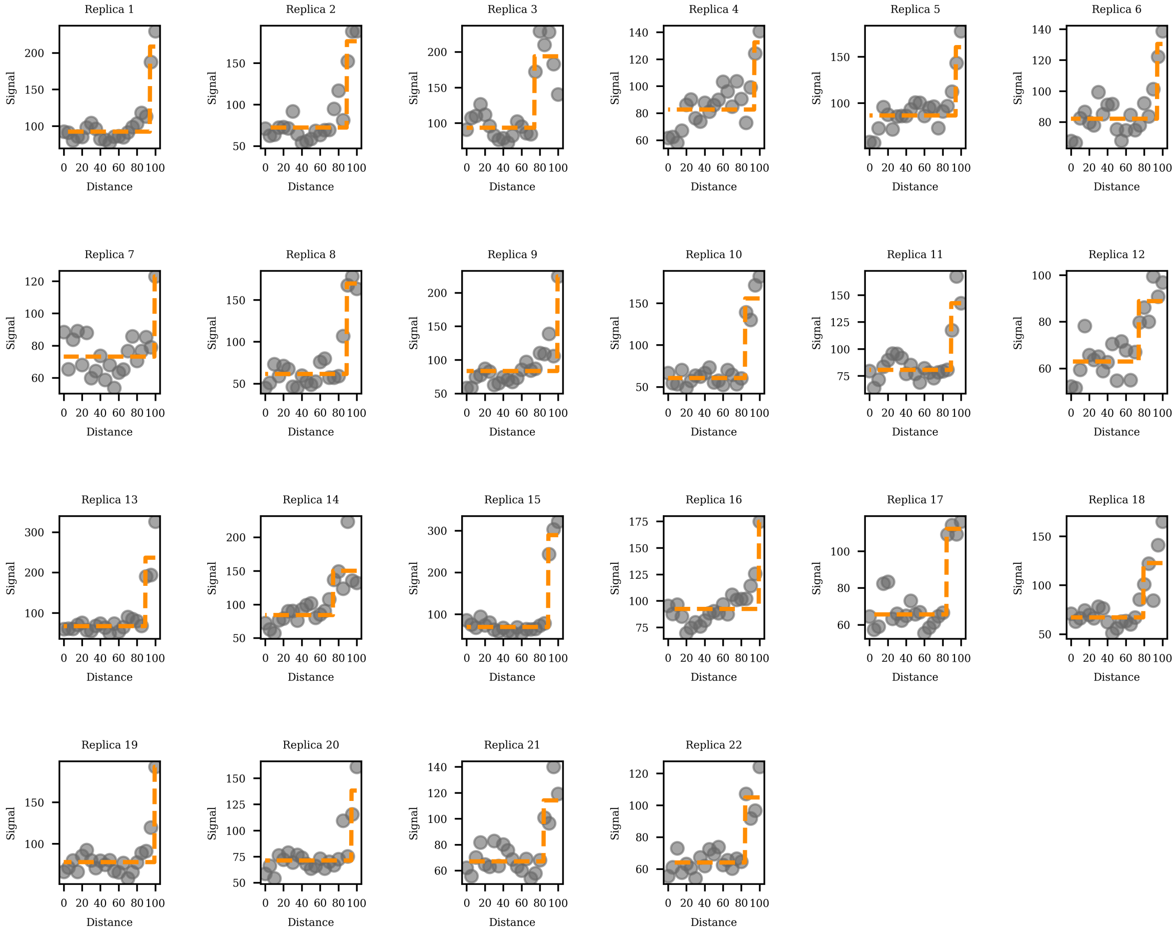

Figure 9: Individual fittings for Shh, steady state.

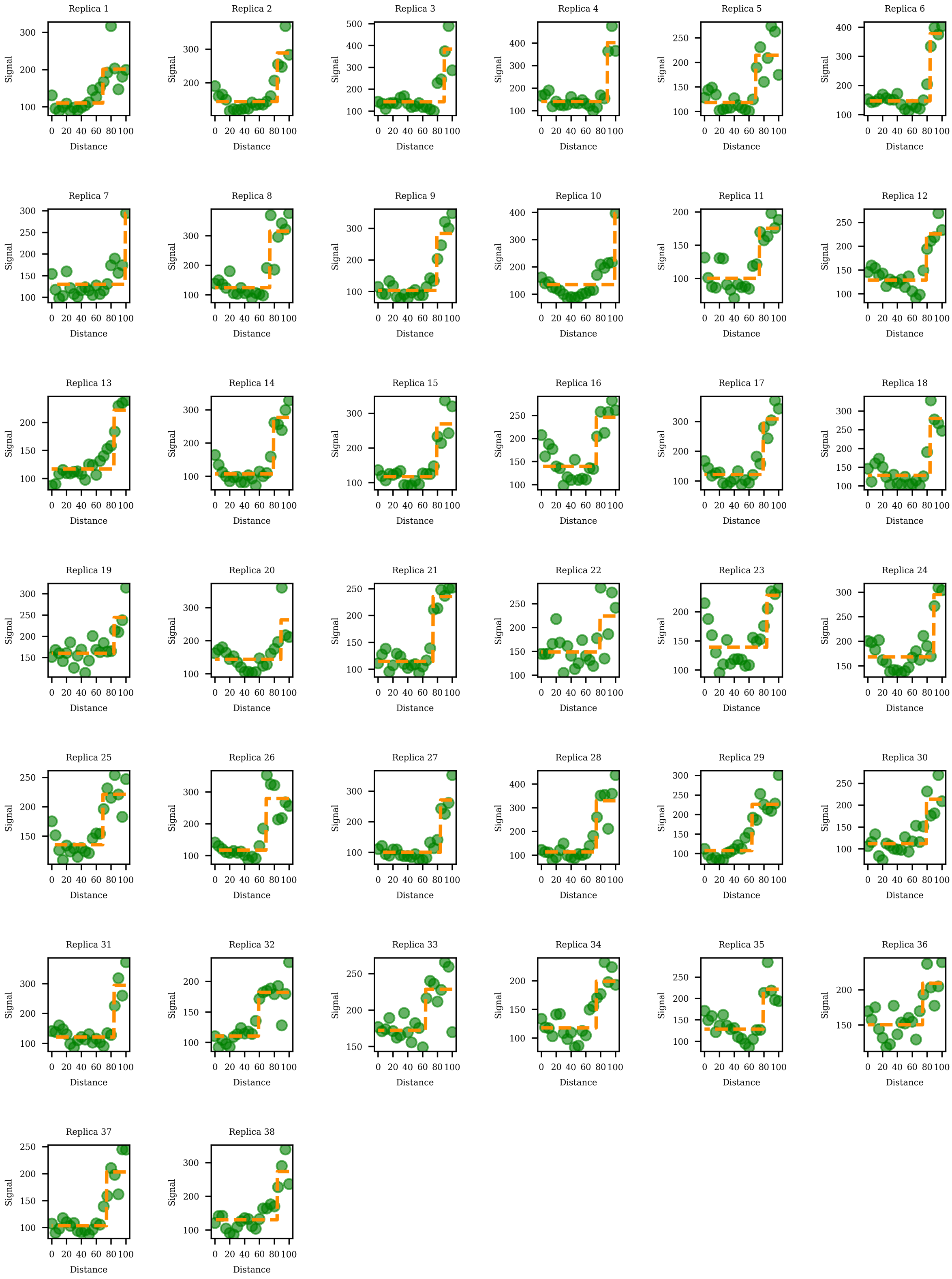

Figure 10: Individual fittings for Shh, 14 dpa.

#### 5.2 Sum of squared errors (SSE)

The figures show the Sum of squared errors (SSE) for different switchpoints in the piecewise constant fitting. Each point represents the SSE calculated for a specific switchpoint (or pair of switchpoints for the three-step function). The switchpoints yielding the lowest SSE are highlighted (orange dot), indicating the best fit for the model.

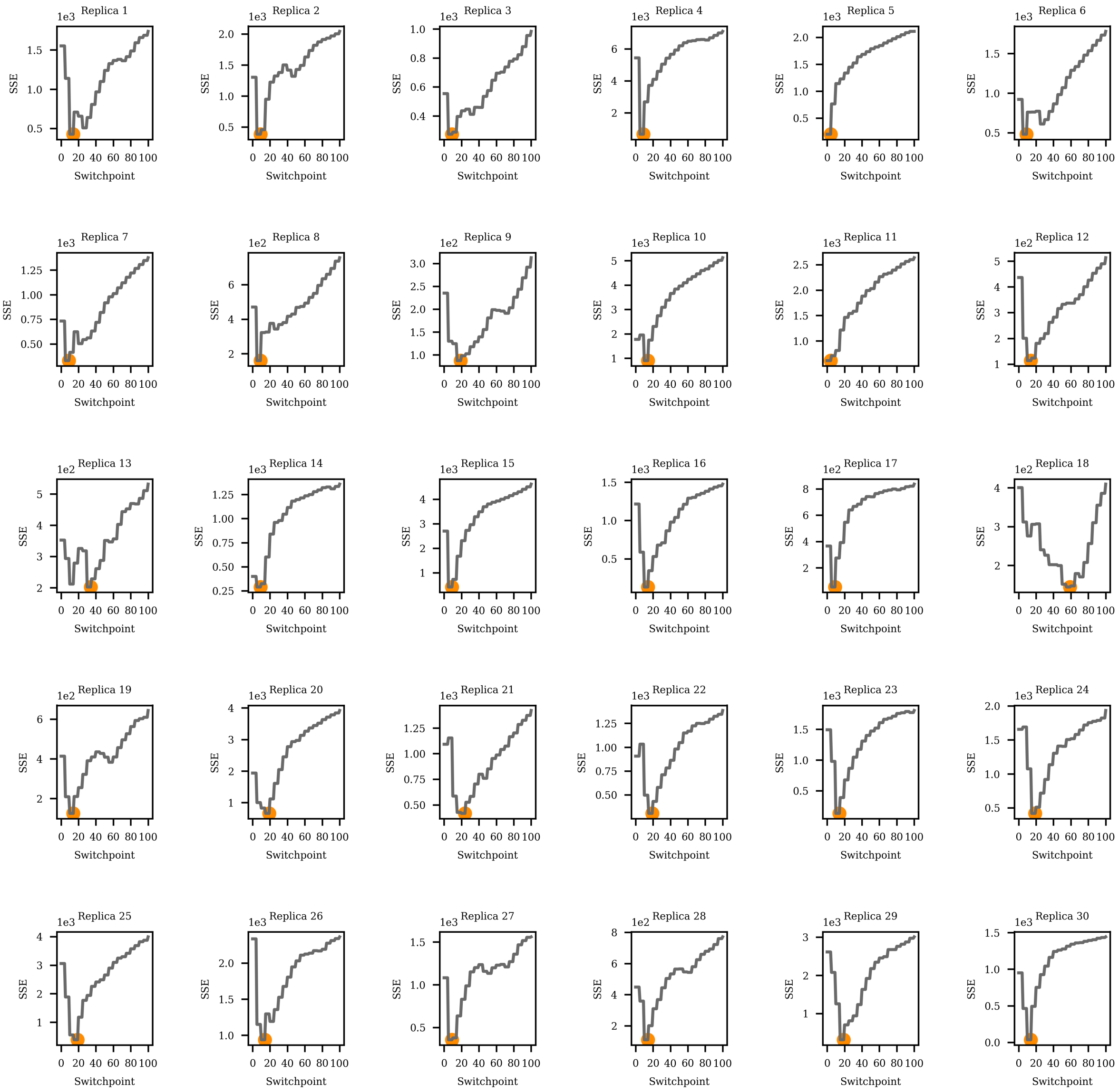

Figure 11: SSE for Msx1, steady state.

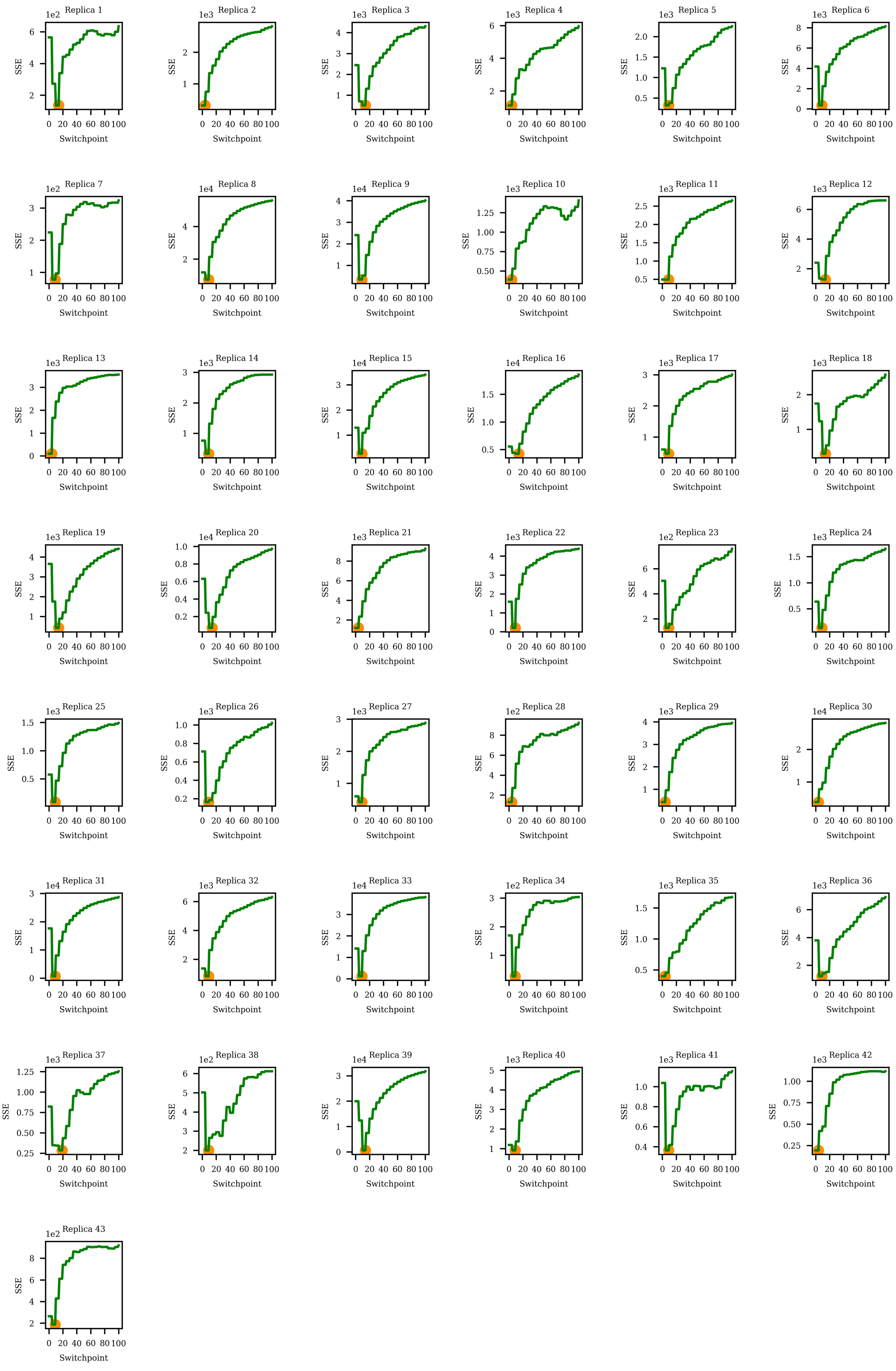

Figure 12: SSE for Msx1, 14 dpa.

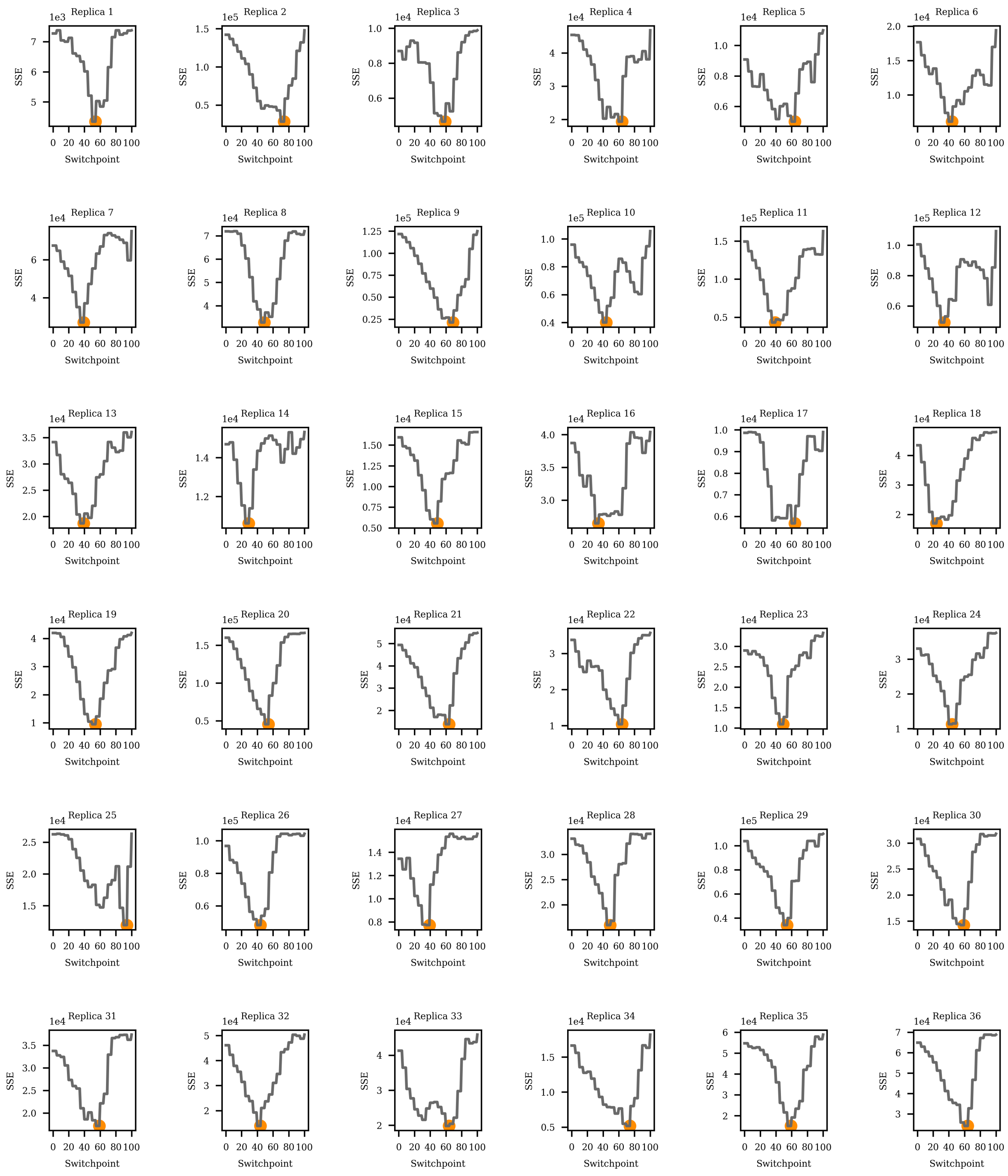

Figure 13: SSE for Nkx6.1, steady state.

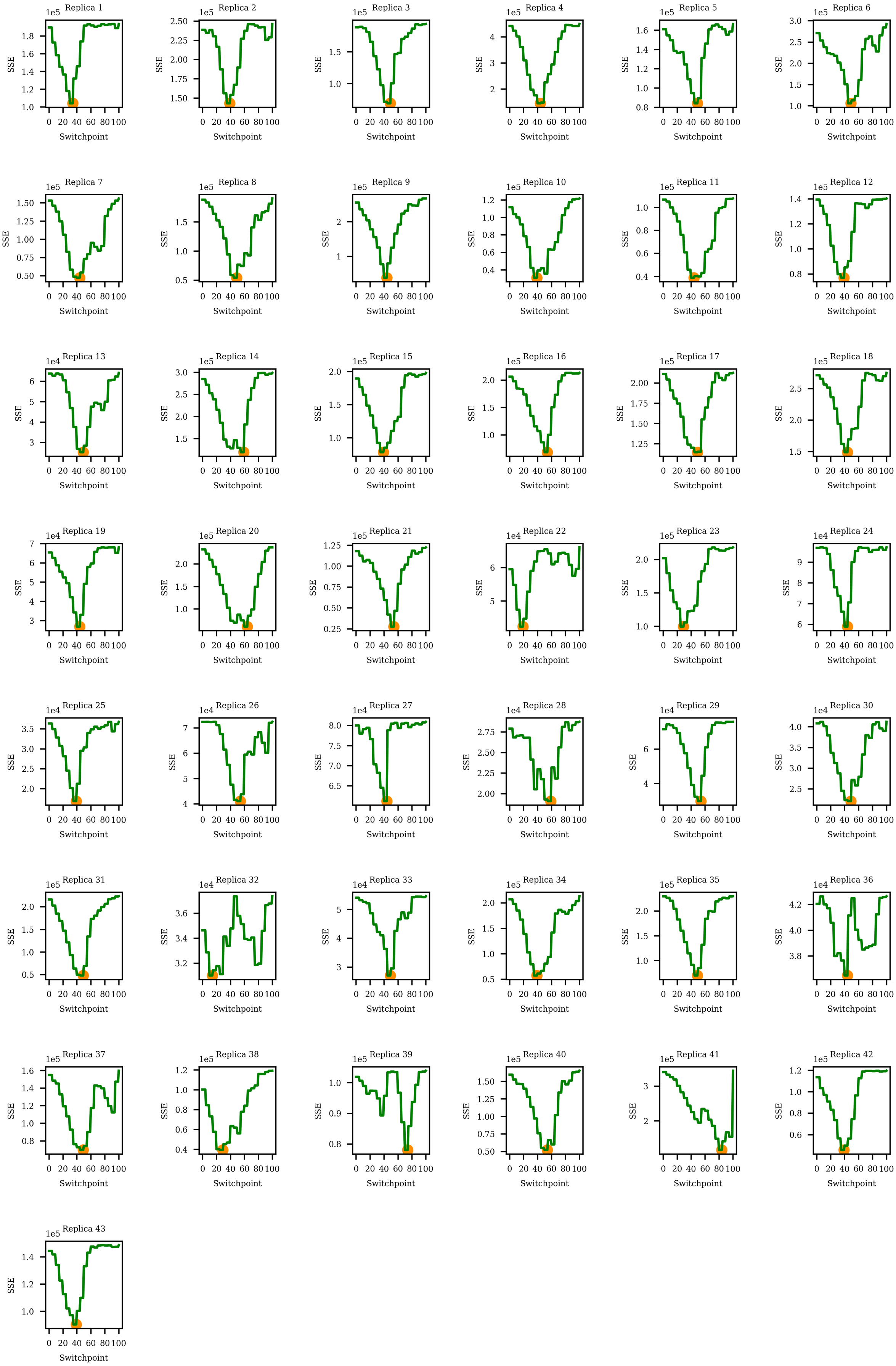

Figure 14: SSE for Nkx6.1, 14 dpa.

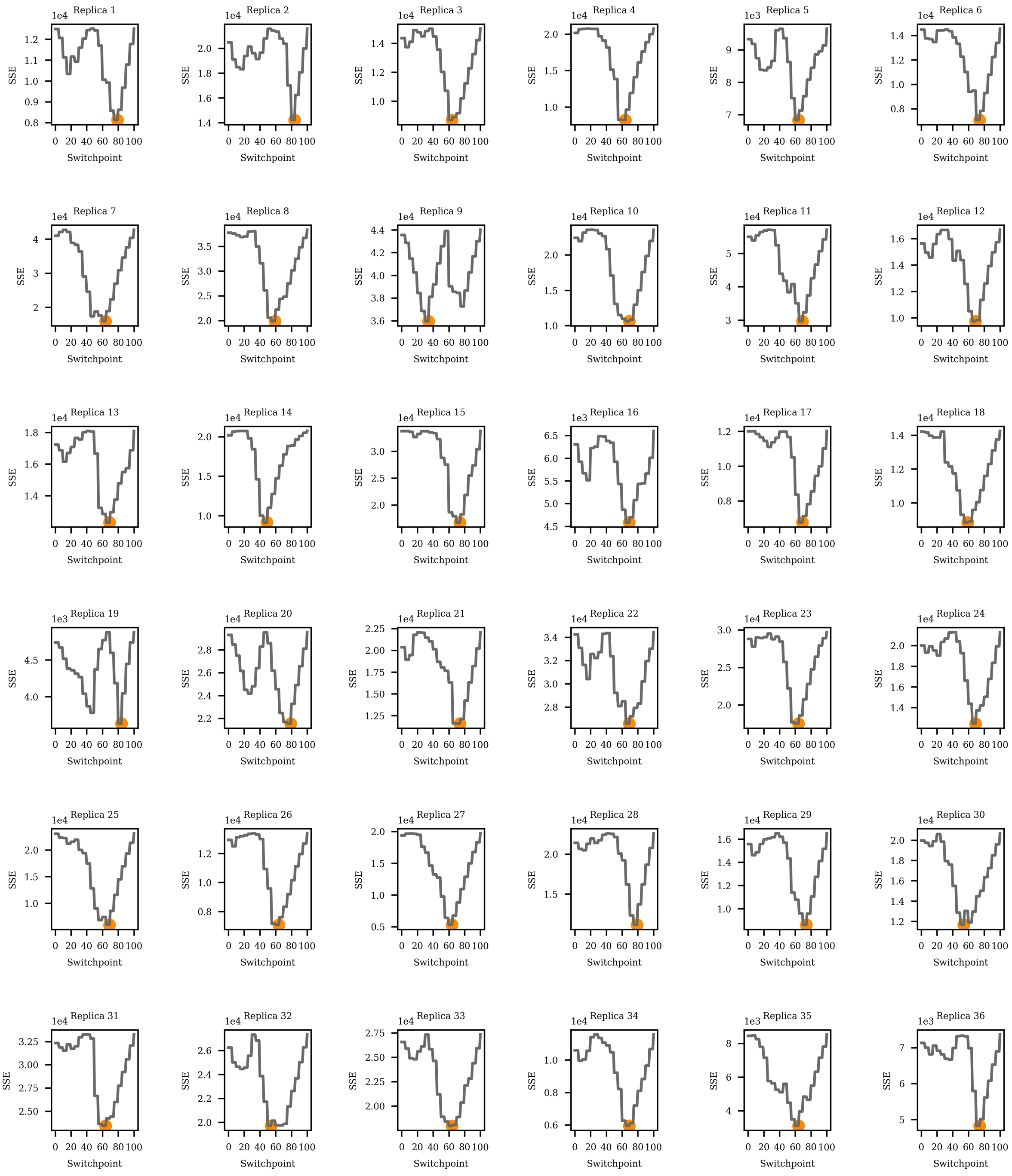

Figure 15: SSE for Pax6, steady state.

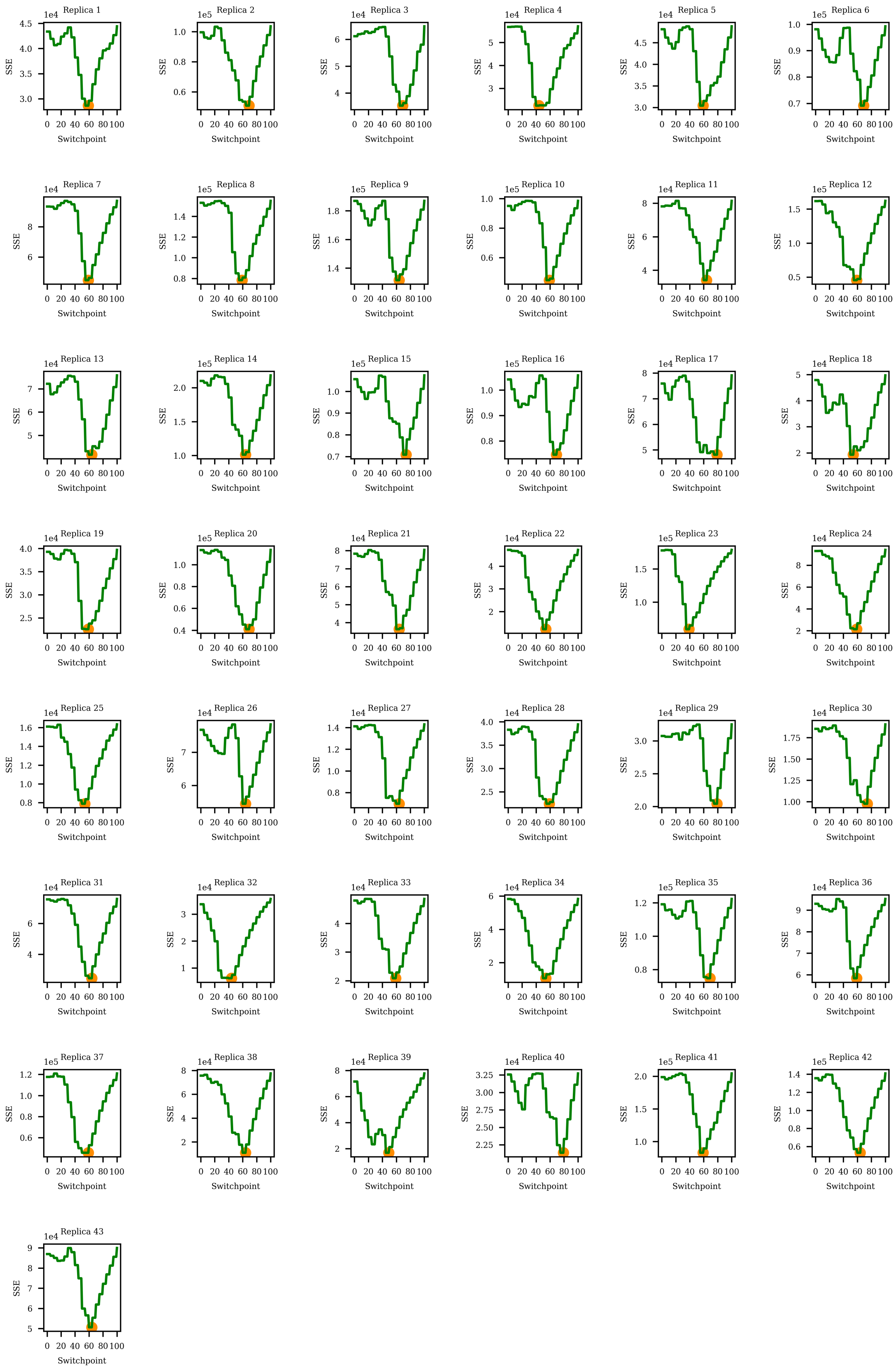

Figure 16: SSE for Pax6, 14 dpa.

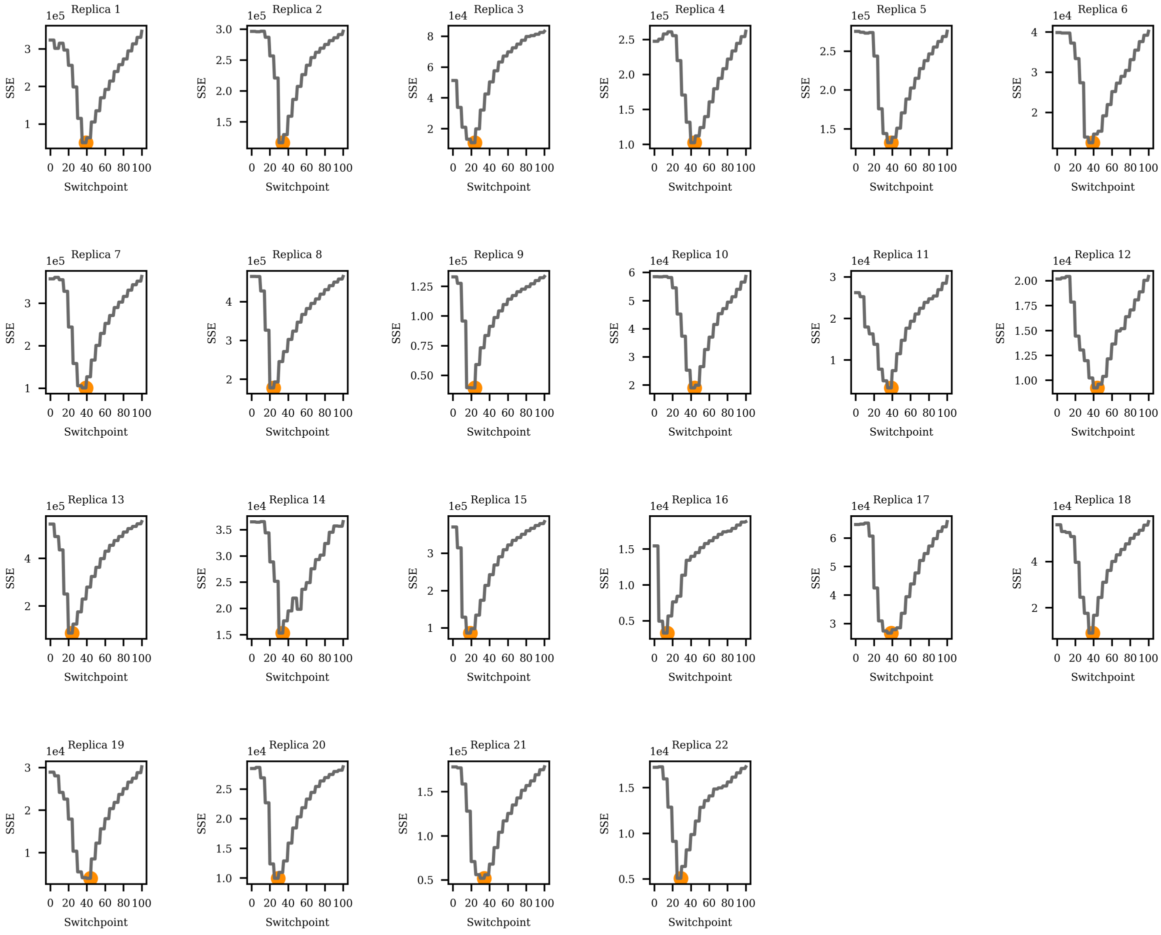

Figure 17: SSE for Pax7, steady state.

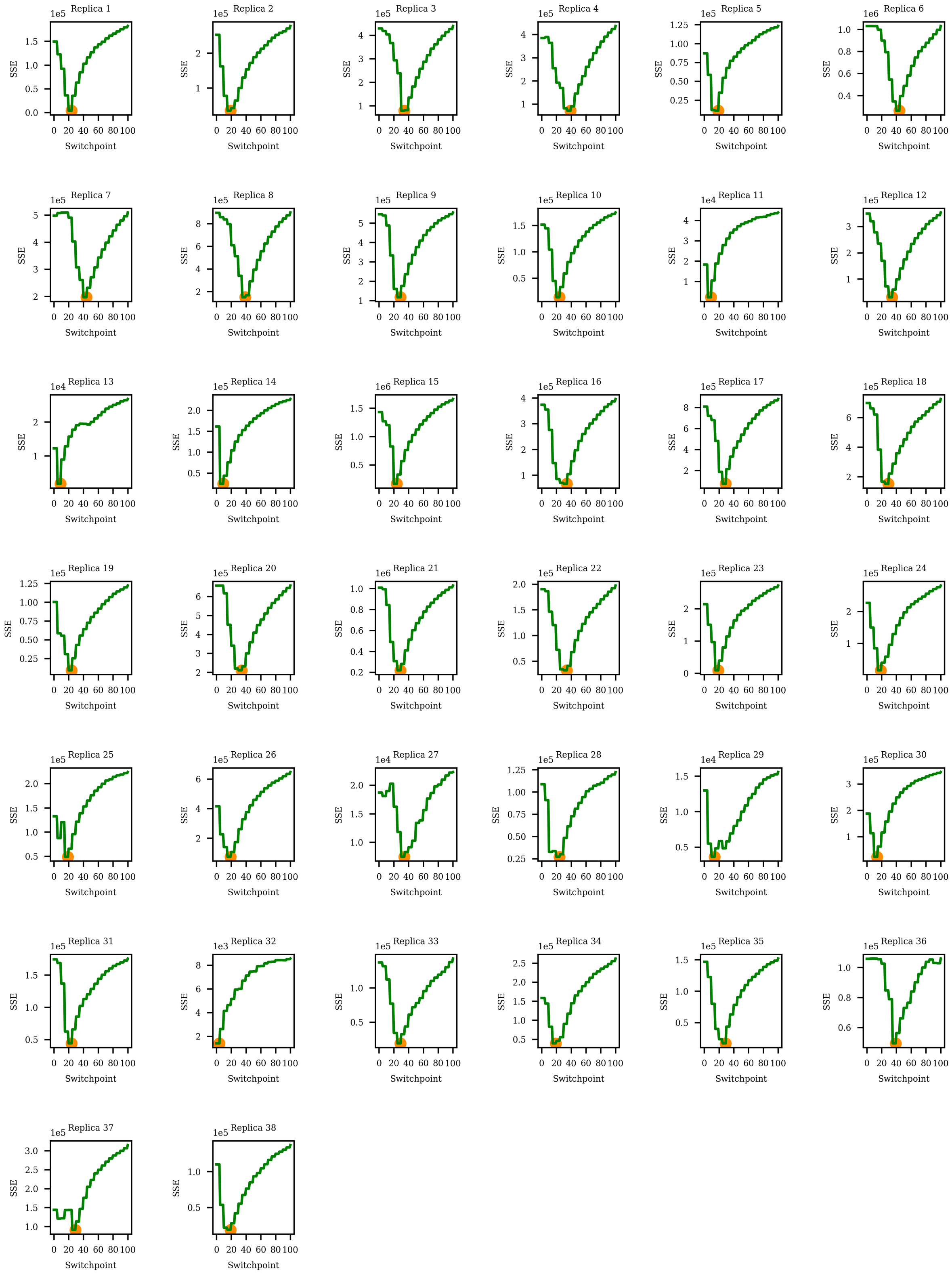

Figure 18: SSE for Pax7, 14 dpa.

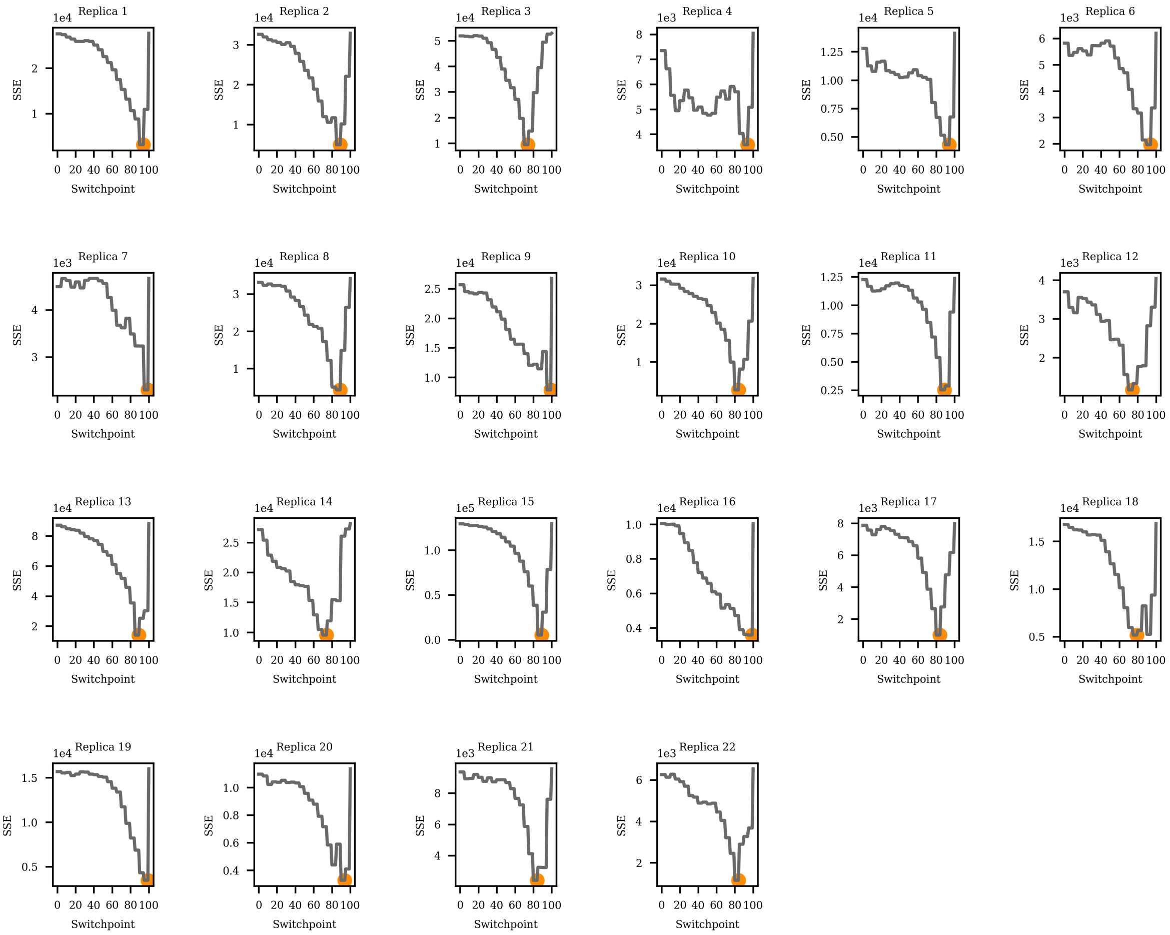

Figure 19: SSE for Shh, steady state.

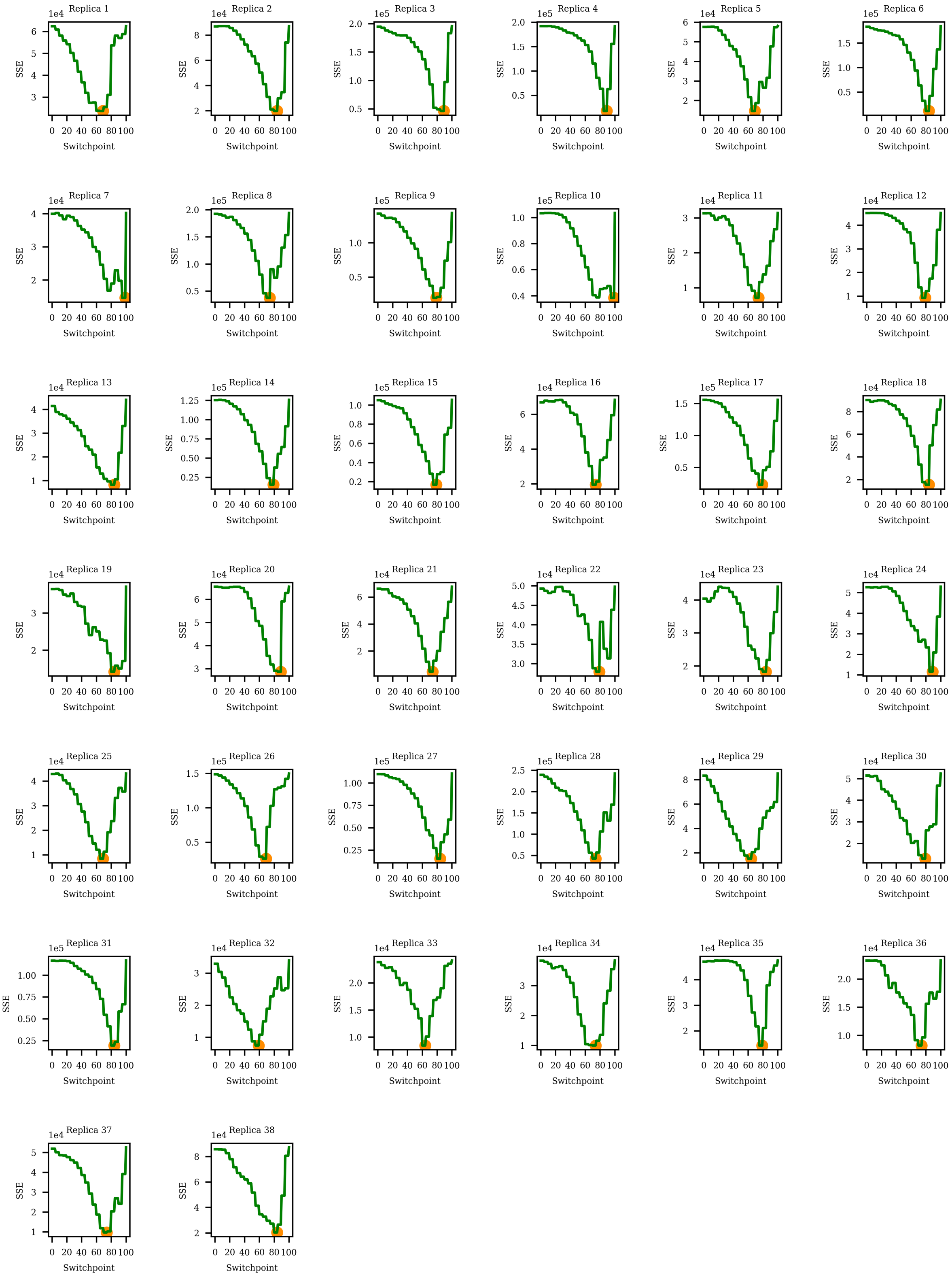

Figure 20: SSE for Shh, 14 dpa.
